# Design and modular assembly of synthetic intramembrane proteolysis receptors for custom gene regulation in therapeutic cells

**DOI:** 10.1101/2021.05.21.445218

**Authors:** Iowis Zhu, Raymond Liu, Axel Hyrenius-Wittsten, Dan I. Piraner, Josef Alavi, Divya V. Israni, Ahmad S. Khalil, Kole T. Roybal

## Abstract

Synthetic biology has established powerful tools to precisely control cell function. Engineering these systems to meet clinical requirements has enormous medical implications. Here, we adopted a clinically driven design process to build receptors for the autonomous control of therapeutic cells. We examined the function of key domains involved in regulated intramembrane proteolysis and showed that systematic modular engineering can generate a class of receptors we call <u>S</u>y<u>N</u>thetic <u>I</u>ntramembrane <u>P</u>roteolysis <u>R</u>eceptors (SNIPRs) that have tunable sensing and transcriptional response abilities. We demonstrate the potential transformative utility of the receptor platform by engineering human primary T cells for multi-antigen recognition and production of dosed, bioactive payloads relevant to the treatment of disease. Our design framework enables the development of fully humanized and customizable transcriptional receptors for the programming of therapeutic cells suitable for clinical translation.

## INTRODUCTION

Cellular function is influenced by both external and internal stimuli, with responses to these stimuli encoded in the genome. Having control over the cellular transcriptional response to a defined external stimulus allows for the development of living, cell-based therapies with programmed therapeutic functions beyond the natural capabilities of a cell. In pursuit of this goal, several synthetic receptor platforms have been developed, including the Tango (Barnea et al., 2008; Kroeze et al., 2015) and Modular Extracellular Signaling Architecture (MESA) (Daringer et al., 2014) systems, as well as the synthetic Notch receptor (synNotch) (Morsut et al., 2016). Notch and synNotch are type 1 transmembrane proteins that activate through regulated intramembrane proteolysis (RIP), a sequential process that involves ADAM protease-mediated shedding of the extracellular domain (ECD), γ-secretase-mediated cleavage of the transmembrane domain (TMD), and release of an intracellular transcription factor (TF) that traffics to the nucleus (Morsut et al., 2016; Kopan and Ilagan, 2009; Gordon et al., 2009). SynNotch receptors recognize a user-defined membrane-bound antigen via a high-affinity ligandbinding domain (LBD), such as a single-chain variable fragment (scFv) or nanobody and induce custom gene regulation through release of an engineered TF (Morsut et al., 2016).

The first generation synNotch receptor is a powerful tool for engineering cell circuitry for programmed multicellular morphologies (Toda et al. 2018), localized tumor control (Roybal et al., 2016b, Srivastava et al., 2019), multi-antigen tumor recognition (Roybal et al., 2016a, Williams et al., 2020), and tumor antigen density discrimination (Hernandez-Lopez et al., 2021). Engineered receptors thus hold enormous potential for furthering our understanding of basic biological processes and expanding our therapeutic options in treating disease. Translating this preliminary work into human therapeutic applications is therefore an important engineering goal.

Despite its central role in several cell engineering milestones, the original synNotch receptor has known limitations that affect its further technological advancement and potential for clinical translation. These issues include 1) the use of non-human components that could elicit immune rejection, 2) the lack of clear design rules for building well-expressed receptors with a tunable activity profile, and 3) the large size of the receptor and integrated transcriptional circuit. We first observed the overall inflexibility of the original design during our attempts to engineer the human equivalent of synNotch, which is based on murine Notch1 (**Fig. S1A, S1B**). Receptors built using human-derived Notch NRRs resulted in poor activation, high ligand-independent signaling, and/or poor expression (**Fig. S1B, S1C**). Moreover, we discovered that both human and mouse-derived synNotch were incompatible with multiple transcription factors (TFs) beyond the yeast- and herpesvirus-derived Gal4-VP64 (**Fig. S1B**). Motivated by these results, we adopted a systematic approach based on modular protein design to define functional receptor modules, allowing us to re-engineer the synNotch receptor from the ground up (**Fig. 1A**). We show that the receptor extracellular domain (ECD), transmembrane domain (TMD), and intracellular juxtamembrane domain (JMD) have distinct and tunable effects on receptor activity.

**Figure 1.**
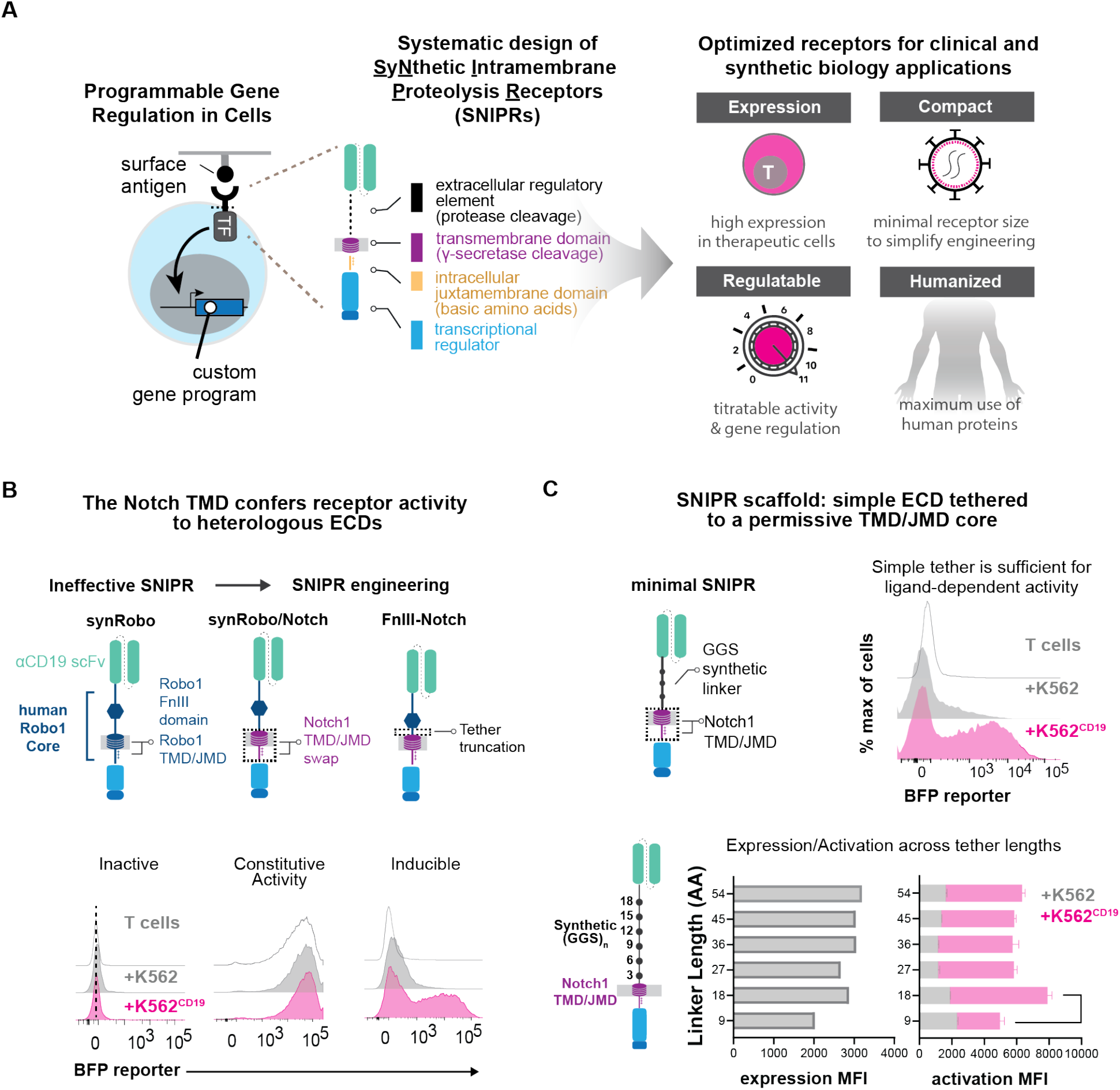
Design of synthetic RIP receptors for customized antigen-dependent gene regulation in therapeutic cells. (**A**) Systematic design of synthetic transcriptional regulatory receptors. Receptors are comprised of a LBD, an extracellular domain (ECD), a transmembrane domain (TMD), a juxtamembrane domain (JMD), and a transcription factor (TF). Receptor circuits are designed to maximize human components, minimize size, express highly in therapeutic cells, and deliver a regulatable level of a therapeutic in response to surface-bound ligand at disease sites **(B)**. A synRobo receptor replaces the Notch1 core with one from human Robo1. T cells expressing a CD19 SNIPR-BFP circuit were co-incubated with either K562 or K562^CD19^ sender cells for 24 hours and BFP output was measured using flow cytometry. Compared to synNotch, a synRobo receptor fails to induce BFP. By replacing the TMD and JMD of Robo1 with those of Notch1, control of BFP production is lost. Deletion of a known ADAM10 cleavage site in the Robo1 ECD rescues ligand-dependent receptor behavior. **(C)** Same as B, but with minimal SNIPRs constructed using simple (GGS)_n_ ECDs, and the TMD/JMD from Notch1.

Through this approach, we have systematically designed, assembled, and tested a large family of <u>S</u>yNthetic <u>I</u>ntramembrane <u>P</u>roteolysis <u>R</u>eceptors (SNIPRs). We present critical design principles of synthetic receptors that undergo RIP and showcase a subset of designs within the larger family that have clear advantages for synthetic biology and next-generation T cell therapeutics. These optimized SNIPRs are compact in size, well-expressed, compatible with human and humanized synthetic TFs, readily tunable, and are both highly sensitive and specific to their target ligand. We show that these SNIPRs function robustly in SNIPR-chimeric antigen receptor (CAR) dual antigen-sensing circuits *in vivo*, a therapeutic strategy that enhances tumor specificity and improves therapeutic efficacy of engineered T cells for solid tumors (Hyrenius-Wittsten et al., 2021, Choe et al., 2021). We also show that we can rationally modify SNIPRs to achieve titratable production of therapeutic payloads such as IL-2, enabling spatially controlled and dosed delivery of therapeutic agents by cells specifically to sites of disease. Though we have focused our efforts on T cells, the menu of modular core receptor parts we have characterized can be mined and used for a broad range of applications in synthetic biology, basic biology, and cell therapeutics.

## RESULTS

### SNIPR development through modular assembly of core receptor domains

To engineer SNIPRs, we took a modular approach for receptor assembly to investigate the role of core domains involved in RIP (**Fig. 1A**). The ECD of Notch1 and other RIP family proteins contain regulated sites of ADAM protease-mediated shedding (Brou et al., 2000; Mumm et al., 2000), and ECD mutations can impact this regulation (Gordon et al., 2009). The TMD is the site of γ-secretase-mediated cleavage and release of the intracellular domain into the cytosol (De Strooper et al., 1999). While γ-secretase is believed to cleave a diverse number of peptides (Beel and Sanders, 2008; Haapasalo and Kovacs, 2011), certain TMD mutations are known to negatively impact cleavage efficiency (Huppert et al., 2000). The basic amino acid-rich JMD connects the TMD to the TF, stops translocation of the receptor through the membrane, and interacts with γ-secretase and endocytosis machinery (Le Borgne, 2005).

Through a similar design strategy to that used for synNotch, a prototypical SNIPR, we sought to engineer a second example of a functional SNIPR. We selected human Robo1, a RIP receptor family member known for mediating ligand-directed neuronal pathfinding (Coleman et al., 2010). Like Notch, Robo1 is a type-I transmembrane protein that undergoes ECD shedding upon ligand engagement, followed by γ-secretase-mediated TMD cleavage to free the cytoplasmic tail (Seki et al., 2010). The putative proteolytic core of Robo1 features an ADAM10 protease-sensitive site that is protected by a compact type III Fibronectin (Fn-III) domain, as well as the receptor TMD and JMD (Coleman et al., 2010). In line with synNotch development, we built a synthetic Robo1 receptor (synRobo) against CD19 that contained the Robo1 proteolytic core and the Gal4-VP64 TF, combining it with the cognate Gal4 DNA response element (RE) controlling a BFP reporter (**Fig. S1A**). Although synRobo expressed at a comparable level to synNotch (**Fig. S1D**), we observed poor reporter activation when T cells expressing the anti-CD19 synRobo were co-incubated with K562^CD19+^ sender cells (**Fig. 1B**). To determine the cause of this stark difference in activity, we substituted two key domains of synRobo, the TMD and JMD, with the equivalent domains from human Notch1. We found that the resulting Robo1/Notch1 chimeric receptor (RoboNotch) was constitutively active, suggesting that the ECD of synRobo was easily shed, but the Robo1 TMD and JMD were not easily processed (**Fig. 1B**). We also found that deletion of the putative ADAM10 protease site in the Robo1 ECD of RoboNotch significantly reduced constitutive signaling and restored ligand-dependent activation. Thus, a canonical protease cleavage site in the ECD is not necessary for receptor function, but the receptor activity remains dependent on ADAM protease activity. (**Fig. 1B, S1E**).

The assembly of a second functional SNIPR through the iterative engineering of parts from Robo1 and Notch1 prompted us to develop a systematic process to explore the principles of receptor design. We thus built a set of SNIPRs to identify critical features of the ECD, TMD, and JMD that are necessary for optimal receptor function. We began with the ECD, constructing a set of SNIPRs with a variable length series of low complexity and flexible glycine-glycine-serine repeats ECDs, an anti-CD19 scFv, and the human Notch1 TMD and JMD. These designs were expressed in human T cells and demonstrated ligand-dependent activation across all tested ECD lengths, as well as a dependence on ADAM protease activity (**Fig. 1C, S1E**). Given that a simple ECD without known protease sites was sufficient for regulated receptor activity, we considered that a broad range of ECDs could be used to assemble functional SNIPRs when paired with a RIP-permissive TMD and JMD. We further hypothesized that additional TMDs and JMDs may be compatible with heterologous ECDs, enabling the modular construction of a customizable family of SNIPRs with diverse activation properties for more customized cellular programming. Importantly, this proposed flexibility in receptor assembly would greatly expand the design space available for engineering clinically compatible receptors.

### ECD engineering controls SNIPR activation parameters

Given our positive results with a Robo-derived ECD and simple linkers, we expanded our survey of ECDs to include synthetic peptides with embedded protease sites, Notch-derived domains, and well-characterized hinge domains sourced from chimeric antigen receptors (CARs). We found that SNIPRs built with synthetic linkers containing exposed ADAM protease sites were constitutively active, while a SNIPR built with a FLAG-tag linker containing an enterokinase cleavage site retained ligand-dependent activity (**Fig. 2A**). Interestingly, an ECD that incorporated fibroblast activation protein (FAP) cleavage sites demonstrated signaling when co-cultured with K562 cells, an effect that was abrogated with the addition of an LBD, suggesting that ECD shedding is dependent on protease availability and cleavage site accessibility (**Fig. 2A, S2A**).

**Figure 2.**
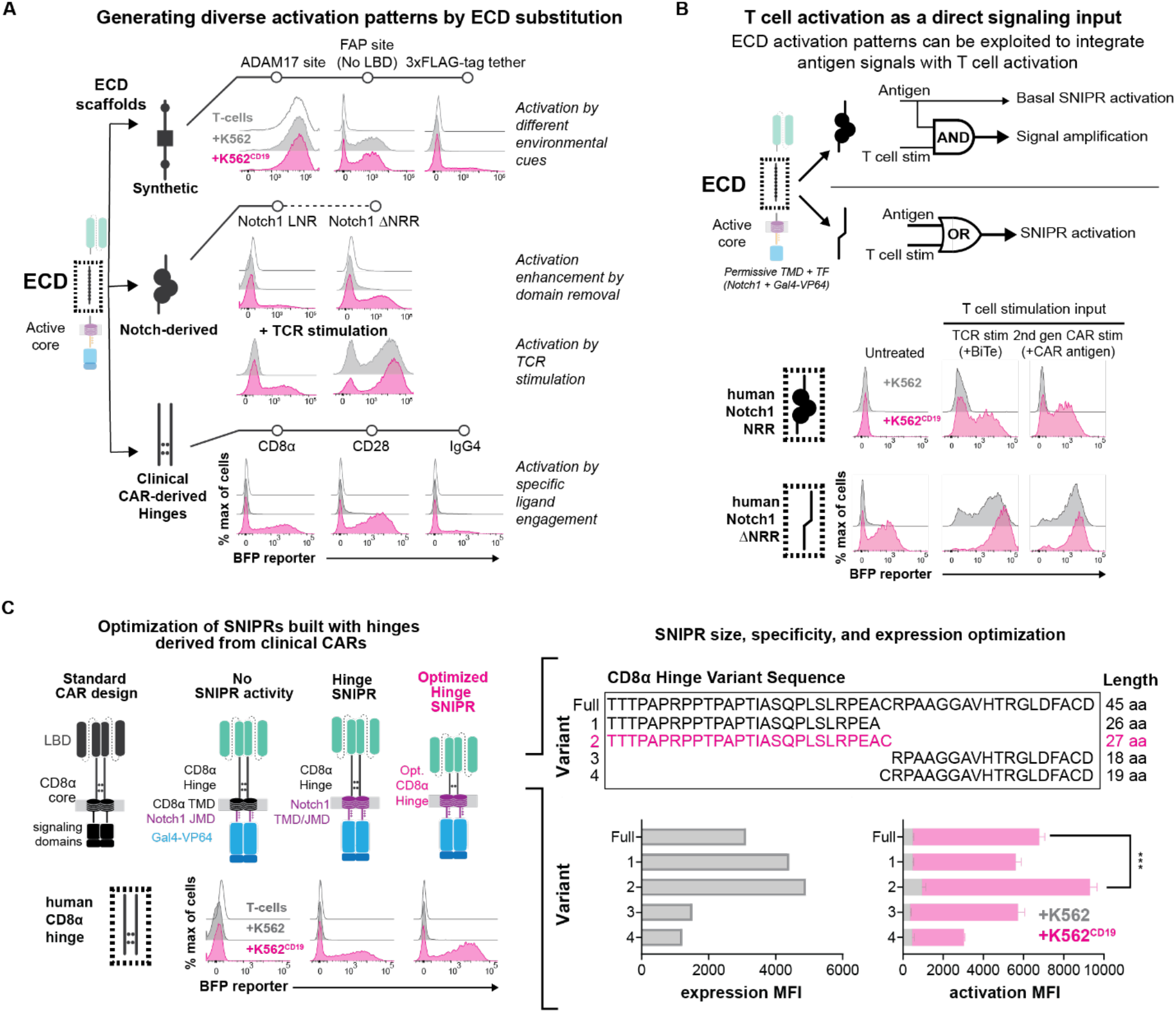
The ECD module defines activation triggers and diversifies sensor functions. **(A)** T cells expressing the indicated anti-CD19 SNIPR-BFP circuit were co-incubated with of either K562 or K562^CD19^ sender cells for 48 hours and BFP output was measured using flow cytometry. SNIPR ECDs with exposed cleavage sites display ligand-independent signaling. Deleting the NRR from synNotch produces a receptor that is sensitive to both ligand and TCR stimulation. A variety of hinge domains utilized in CARs also demonstrate ligand-dependent signaling. **(B)** Same as A, but with two methods of T cell stimulation. A SNIPR with the Notch1 NRR core domain displays enhanced activation with a Bispecific T cell Engager (BiTE) targeting a K562 antigen, and a co-expressed second-generation CAR targeting a separate antigen. A SNIPR with a truncated Notch1 NRR activates with these stimuli independent of the presence of ligand. **(C)** Same as A, but with variations of the CD8α hinge ECD. The CD8α hinge can be optimized to enhance SNIPR expression and activation.

Given the surprising diversity of functional ECDs in SNIPRs, we decided to assess whether the Notch Regulatory Region (NRR), a large majority of the synNotch ECD, was necessary for receptor function. To our surprise, a SNIPR with a full deletion of the NRR (ΔNRR) exhibited strong ligand-induced signaling but was also triggered by T cell activation alone (**Fig. 2A**). This T cell activation-based receptor activity was observed with several methods of T cell activation, including with Bi-specific T cell Engagers (BiTEs) or a co-expressed second generation CAR. In addition, we observed that T cell activation drove the enhanced activation of SNIPRs, such as synNotch, that appeared insensitive to T cell activation alone (**Fig. 2B, S2B**). These data demonstrate that a spectrum of ECDs is compatible with SNIPR construction and that the choice of ECD can significantly impact the fidelity and sensitivity of the resulting receptor (**Fig. 2B, S2B**).

### Clinically oriented ECD engineering

CARs often include a hinge region derived from immunoglobulin-like domains, such as CD8α or CD28, or from more complex trimeric receptors (e.g., OX-40) in the ECD that affects the oligomeric state, flexibility, and general ligand-binding properties of the receptor (Guedan et al., 2019). We found that, when used in our SNIPR designs, CD8α and CD28-based hinge ECDs exhibited high expression and receptor activation, with the CD8α hinge exhibiting reduced ligand-independent signaling (**Fig. 2A, S2C**). However, the CD8α hinge displayed ligand-independent signaling with T cell activation, especially in CD8+ T cells (**Fig. S2C, S2D**). Given these results, we devised a strategy to improve the functionality of the CD8α hinge ECD through a series of N-terminal and C-terminal truncations. From testing four truncation variants, we found that the 27 amino acid N-terminal region of the CD8α hinge displayed enhanced expression and minimal ligand-independent activity with T cell activation (**Fig. 2C, S2E**). This optimized CD8α hinge SNIPR was nominated for additional development due to its efficient, high-fidelity activation and compact size, with the full optimized anti-CD19 CD8α hinge SNIPR with Gal4-VP64 being only 1.65kB in length whereas the original synNotch is 2.45kB, a 33% reduction in size.

### TMD and JMD engineering can tune receptor activity

The Notch1 TMD and JMD are functional with a broad set of ECDs (**Fig. 2A**). We next investigated the characteristics of the domains that make them functional, and whether additional TMD or JMD sequences were compatible with SNIPR assembly. To do this, we compiled a list of proteins known to undergo RIP and extracted their TMD and JMD sequences (Haapasalo and Kovacs, 2011) (**Fig. 3A, Supplemental Table 1**). For this study, we defined the JMD as a stretch of basic amino acids (R/K/H) beginning immediately C-terminal to the TMD and ending before three consecutive non-basic amino acids. Due to the diversity observed in both TMDs and JMDs, we decoupled the TMD-JMD pair into two separate modules for screening and compiled two libraries, one of 88 TMDs and another of 76 JMDs. We then inserted them individually into a human synNotch scaffold, replacing the respective Notch1 components, and screened them using a Jurkat reporter cell line in an arrayed format (**Fig. 3A**). We quantified activity, analyzed the TMDs displaying >50% of Notch1 TMD activity for sequence similarities, and identified additional TMDs and JMDs of interest for further testing (**Fig. 3B, 3C)**.

**Figure 3.**
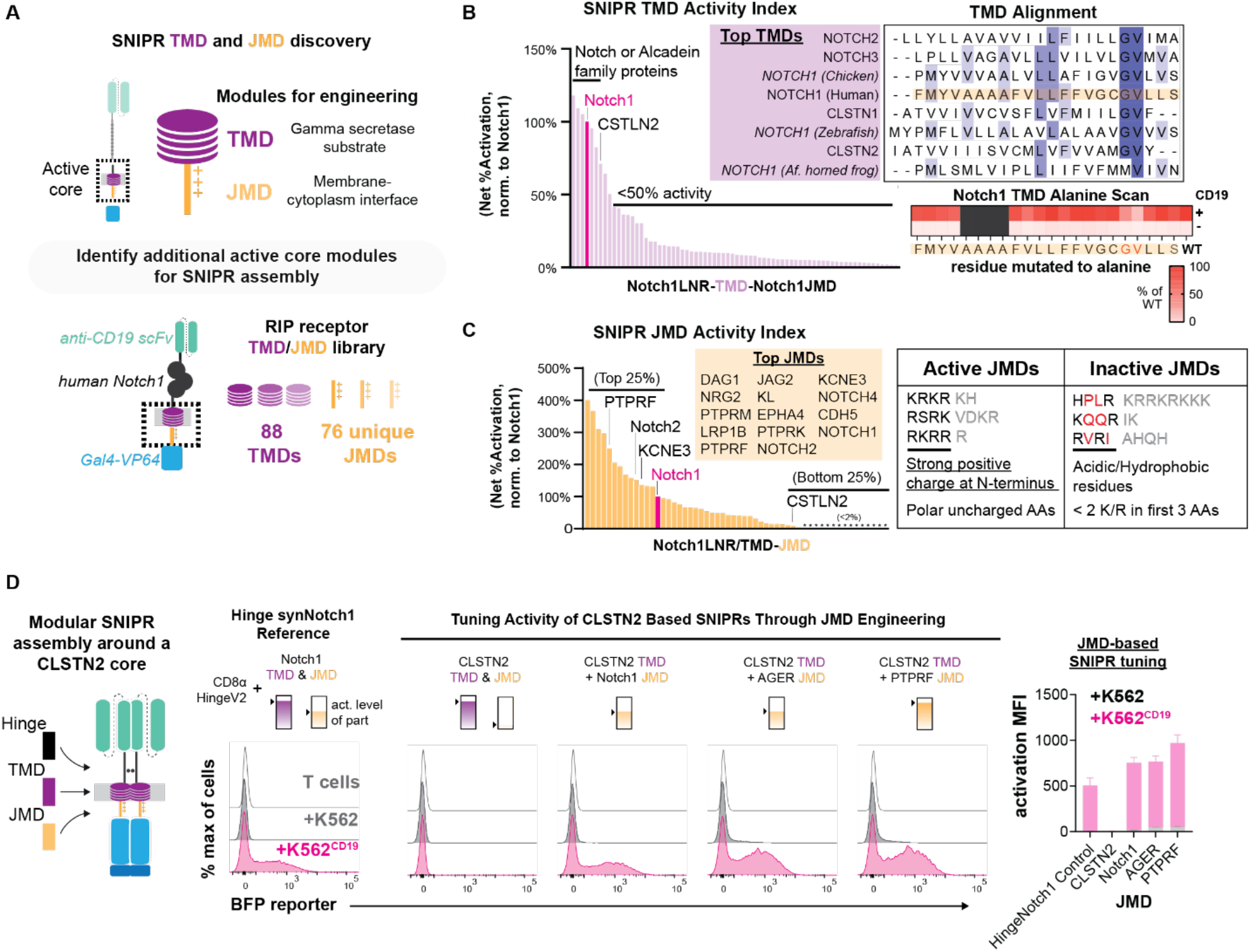
Transmembrane and juxtamembrane domain libraries enable modular assembly of novel SNIPR architectures. **(A)** To identify functional receptor TMDs and JMDs for modular assembly, 88 TMDs and 76 JMDs were cloned into a human synNotch scaffold, replacing either the Notch1 TMD or JMD, respectively. Jurkat T cells expressing an inducible BFP reporter were transduced with these SNIPR libraries in an arrayed format. **(B)** Jurkat T cells were co-incubated with K562 or K562^CD19^ sender cells for 24 hours and BFP output was measured using flow cytometry. Net %BFP+ activation was calculated by subtracting %BFP_K562_ from %BFP_CD19_ and was normalized to the human Notch1 TMD. An alignment of the best performing TMDs shows a common Gly-Val motif (Dark blue = >80% agreement with consensus sequence, blue = >60% agreement, light blue = >40% agreement). An alanine scan of the human Notch1 TMD in primary T cells supports the importance of this motif. **(C)** Same as B, but with the JMD library. High-performing JMDs are strongly basic at their N-termini and may include polar residues but not acidic or hydrophobic residues. **(D)** Compared to a reference SNIPR containing the Notch1 TMD/JMD, a SNIPR containing the CLSTN2 TMD/JMD (CLSTN2) is inactive, but receptor function is restored when the CLSTN2 JMD is replaced with the Notch1, AGER or PTPRF JMD.

From our TMD screen, we discovered that the top performing TMDs were mainly from the Notch and CLSTN protein families, with the activity of most TMDs below 50% of that of Notch1 (**Fig. 3B**). Alignment of the Notch and CLSTN TMD sequences reveals a common c-terminal glycine-valine motif associated with γ-secretase cleavage. Previous studies have shown that these sites are essential for efficient intramembrane processing by presenilin (Vooijs et al., 2004; Okochi, 2002). To determine the importance of this motif in SNIPR signaling, we performed an alanine scan within the Notch1 TMD in primary human T cells, using the optimized CD8α hinge Notch ECD and Notch1 JMD (**Fig. 3B**). Although receptor expression was not reduced (**Fig. S3A**), we found that substitution of the glycine (G318A) and invariant valine (V319A) reduced receptor activity by 47% and 75%, respectively. Background signaling activity from these receptor variants was also lower, consistent with decreased processing as seen in other studies examining this proteolytic site (**Fig. 3B, S3B**) (Vooijs et al., 2004; Okochi, 2002). In addition, two otherwise non-functional TMDs, from Robo1 and AGER, could be made functional through the addition of a Gly-Val motif and removal of bulky residues near the TMD c-terminus (**Fig. S3C**).

In contrast to the TMD screen, our results from the JMD screen showed that JMDs sourced from a diverse set of proteins were effective in a SNIPR context (**Fig. 3C, S3D**). We found that the top JMD sequences favored highly basic residues immediately adjacent to the membrane, with basic or polar residues composing the first 4 to 6 amino acids, and at least two R/Ks within the first 3 amino acids. Hydrophobic or acidic residues within this stretch were found to severely inhibit receptor activation. Replacement of either the Notch1 TMD or JMD did not affect SNIPR sensitivity to T cell activation alone, suggesting that T cell activation affects SNIPR activity at the level of ECD cleavage.

### Building non-Notch SNIPRs from a set of functional parts

Thus far, all functional SNIPRs we have studied include sequences derived from Notch family members. To demonstrate the versatility of our modular assembly approach, we proceeded to engineer a functional ligand-activated receptor without Notch domains, combining the optimized CD8α hinge ECD with the CLSTN2 TMD and functional JMD modules discovered in our screens. Although all receptors expressed (**Fig. S3E**), the otherwise-active CLSTN2 TMD did not function with its cognate JMD (RVRIAHQH), an expected result given the poor performance of the CLSTN2 JMD in our JMD screen. However, receptor functionality was restored by replacing the JMD with that of AGER (RRQRR) or PTPRF (KRKRTH), two potent JMDs identified in our screen (**Fig. 3D, S3E**). Our ability to build new functional SNIPRs from a set of functional parts demonstrates that our approach to receptor assembly can generate receptors that can be easily modified to tune receptor sensing and activity (**Fig. S3E**).

### Precision control and customization of T cell therapeutics with SNIPRs

Based on the design principles of SNIPRs we uncovered, we next assembled receptors from a menu of ECD, TMDs, and JMDs with a range of activation characteristics. Our design criteria included robust expression, a range of ligand-dependent activation levels, and low ligand-independent activation under both basal and activated T cell conditions (**Fig. 4A, S4A**). For the ECD, we selected the optimized CD8α hinge, due to its strong expression, compact size, and selective response to ligand. We then screened through a selection of high-performing TMDs and JMDs from our screens, using a constant Notch1 JMD or TMD, respectively, for simplicity. From this process, we decided to keep the Notch1 TMD due to its robust activation and best-in-class levels of ligand-independent signaling, along with the ready availability of mutants for tunability (**Fig. 4A**). We screened this ECD-TMD combination against a panel of JMDs, choosing a set of SNIPRs with a range of activation levels. Although SNIPR expression levels varied between TMDs and JMDs, these differences did not correlate perfectly with SNIPR activation, supporting a role for the JMD in affecting activity beyond impacting cell surface expression. The assembled set of SNIPRs remained sensitive to ADAM protease and γ-secretase inhibition, suggesting a continued role for these proteases in SNIPR activation (**Fig. S4B**). They are also compatible with a variety of ligand-binding domains (**Fig. 4B, S4G**).

**Figure 4.**
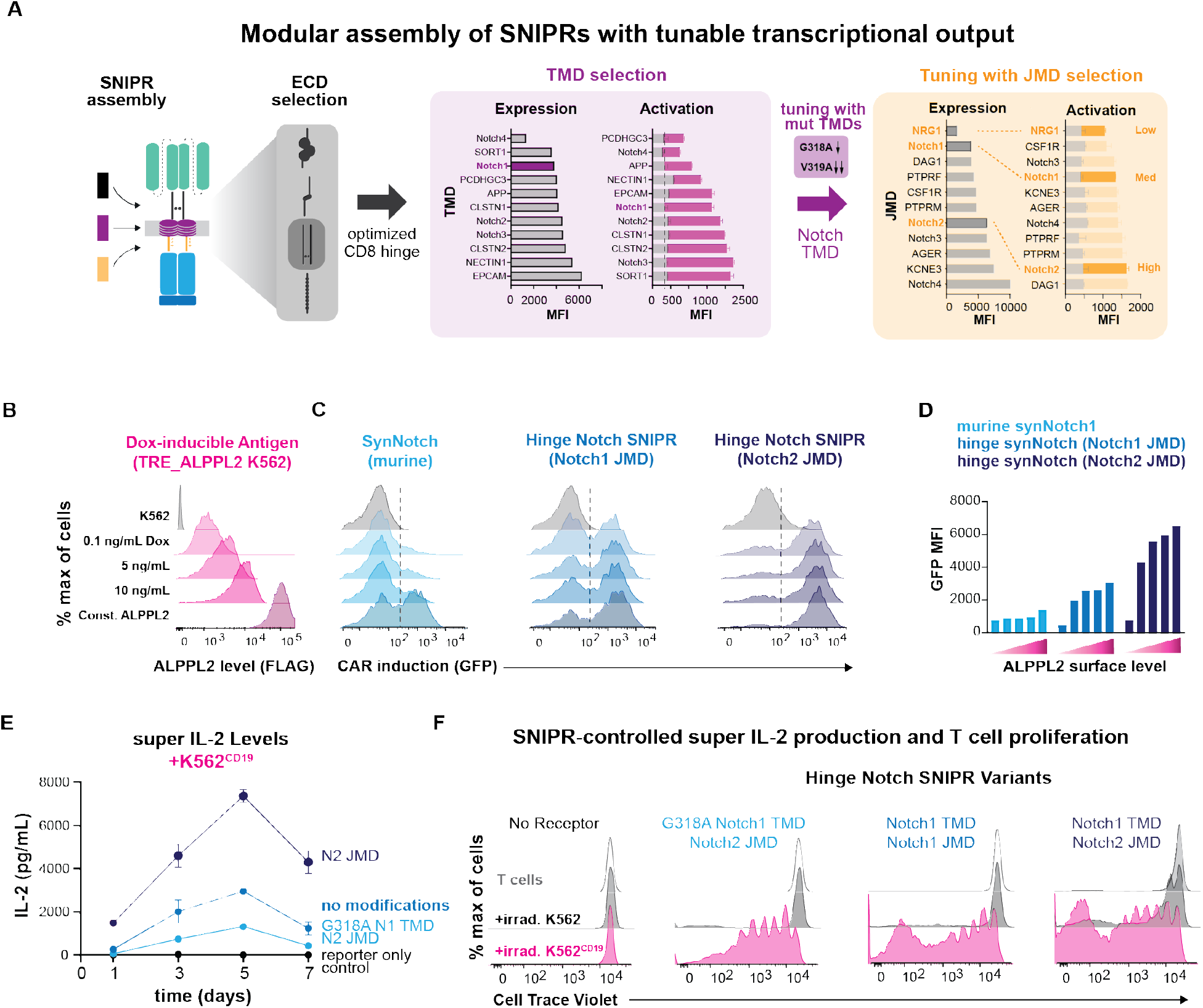
Enhanced sensitivity and tunable gene regulation through SNIPR engineering. **(A)** T cells expressing high-performing SNIPR-BFP circuits were co-incubated with sender cells for 48 hours. BFP output was measured using flow cytometry. The Notch1 TMD was selected for further testing. Three JMDs and two TMD alanine mutants were selected to produce a wide output range. **(B)** K562 cells transduced with a doxycycline-inducible FLAG-tagged ALPPL2 cassette express ALPPL2 in a dose-dependent manner. **(C)** CD4+ T cells expressing anti-ALPPL2 SNIPR-MCAM CAR circuits were co-incubated with sender cells for 48 hours and CAR output was measured using a t2a GFP system. **(D)** Graphical representation of C. **(E)** CD4+ T cells expressing anti-CD19 SNIPR-super IL-2 circuits were co-incubated with irradiated sender K562 cells in media without IL-2. Supernatant IL-2 concentration was assayed using ELISA. **(F)** T cells stained with Cell Trace Violet were co-incubated with irradiated sender cells in media without IL-2 for 9 days. T cell proliferation was measured using flow cytometry.

Having extensively investigated the range of domains that can be used to build functional SNIPRs, we next determined how to control the therapeutic function of engineered cells through our novel receptors. Many cancers adapt to CAR T cell therapy through antigen escape, downregulating their levels of surface CAR antigen (Majzner and Mackall, 2018). Having observed that our new SNIPRs exhibited improved activation to CD19, we decided to test their ability to sense low surface antigen levels. To do this, we activated T cells engineered with ALPPL2 targeted SNIPRs with a K562 sender cell line transduced to induce expression of the tumor-specific antigen ALPPL2 in response to different levels of doxycycline (Hyrenius-Wittsten, 2021) (**Fig. 4B, S4C, S4D**). Using this system, we demonstrated that the optimized CD8α Hinge Notch SNIPR is more sensitive to low ligand levels than synNotch with no increase in basal activity and that use of the Notch2 JMD further boosts sensitivity (**Fig. 4C, 4D**). These data suggest that SNIPRs could be useful in a wider array of immunotherapeutic applications where antigen density is low or heterogenous across the tumor mass.

Immune cell function is regulated by cytokines in a dose-dependent fashion, and serious side effects occur when a high dose of cytokines is given systemically as an immunotherapeutic (Pachella et al., 2015). Given that SNIPR activity is readily tuned through the TMD and JMD, we wanted to showcase how SNIPRs can be used to drive defined levels of the immunotherapeutic and T cell growth factor, IL-2. To do this, we built single viral vector constructs containing SNIPRs with a range of activity levels and an inducible super IL-2 cassette (Levin et al., 2012) (**Fig. S4E**). Using ELISA, we observed that CD4+ T cells expressing a SNIPR with the enhancing Notch2 JMD modification secreted higher amounts of IL-2 into the supernatant in response to ligand expressed on K562 sender cells that had been irradiated to prevent culture overgrowth, while those with the additional dampening TMD mutation G318A secreted lower amounts of IL-2 (**Fig. 4E**). The different amounts of induced super IL-2 produced by each SNIPR circuit correlated with T-cell proliferation rates, exhibiting our ability to tune therapeutic T cell activity, and did not correlate purely with SNIPR expression levels (**Fig. 4F, S4F**). While we use the example of IL-2 to demonstrate the novel capabilities of the SNIPR platform, this principle of receptor tuning can be applied toward a broad range of therapeutic programs (Roybal et al., 2016a).

### Development of fully humanized SNIPRs with potential for clinical translation

We have systematically explored the core regulatory domains (ECD, TMD, and JMD) that control the ligand-dependent cleavage of SNIPRs and have identified optimized cores. However, the potent synthetic Gal4-VP64 TF is a potential design liability for clinical translation as it is derived from yeast (Gal4) and herpesvirus proteins (VP64). To engineer a fully humanized receptor, we constructed TFs comprised of DNA-binding domains (DBD) fused to the transactivation domain of human NF-κB p65. We examined both DBDs sourced from human proteins, as well as engineered orthogonal synthetic zinc fingers (synTFs), for their ability to function in the SNIPR context (**Fig. 5A**). Human protein-derived DBDs are advantageous for minimizing immunogenicity, whereas synTFs minimize off-target effects as verified by RNAseq (Israni et al., 2021). Human protein-derived DBDs were chosen based on size and lack of expression in T cells (Uhlen et al., 2015), and included the eye development-associated paired box protein Pax-6 (Pax6) (Xu et al., 1999) and the liver-specific protein hepatocyte-nuclear factor 1-alpha (HNF1A) (Roscilli et al., 2002). SynTF candidates were selected for their orthogonality and potent transcriptional activity (Israni et al., 2021). RE cassettes for these TFs were constructed by tandem assembly of cognate binding motifs upstream to a minimal promoter. These RE cassettes proved to be orthogonal in T cells, as they were not activated in the absence of target cells (**Fig. 5B**). Humanized SNIPRs activated in the presence of target cells, and activation varied across TFs, suggesting that circuit function is subject to the efficiency of each TF in driving transcriptional activation (**Fig. 5B**). To examine whether TF compatibility extends to the original synNotch receptor, we tested the two TFs with the highest MFI of activation, HNF1A and ZFN10, with the mouse synNotch and humanized receptor variant, and found that neither expressed nor activated as efficiently as the equivalent SNIPR utilizing the optimized CD8α hinge (**Fig. 5C, S5A**). We also examined whether a fully humanized anti-CD19 SNIPR can eliminate target cells via BCMA-CAR payload induction. We found that HNF1A-based receptor circuits induced BCMA-CAR expression at a slower rate as compared to Gal4-VP64, but at sufficient levels to clear *in vitro* tumor targets (**Fig. 5D**). These data show that the enhanced features of optimized SNIPR design enables compatibility with a broader range of TFs.

**Figure 5.**
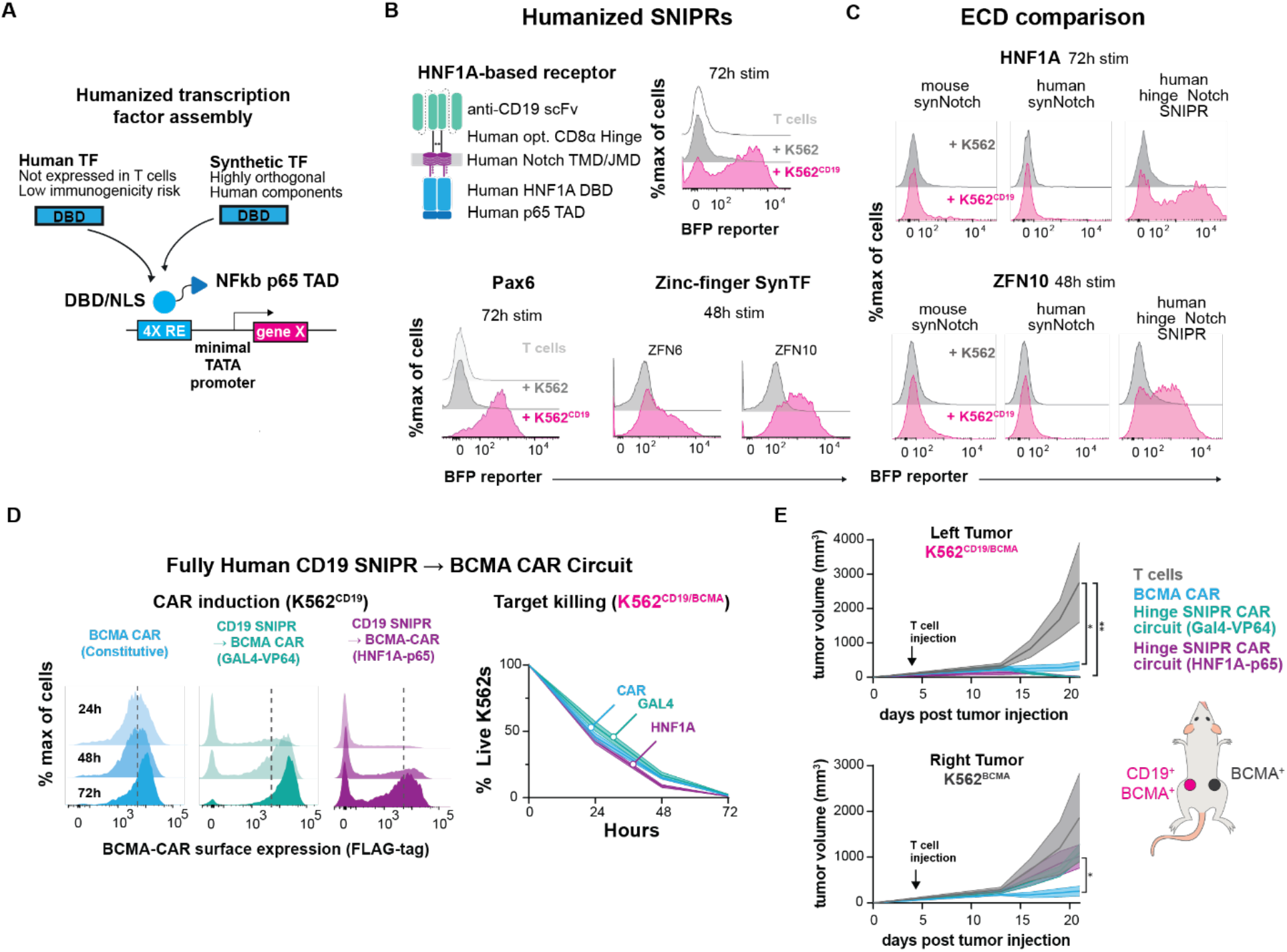
Humanization of SNIPRs to reduce immunogenicity potential for cell-based therapies. **(A)** Human transcription factor engineering. Humanized TFs were constructed by fusing the DNA-binding domain of human TFs not expressed in T cells, or engineered orthogonal synthetic zinc-finger TFs, to the NF-κB p65 transactivation domain. A cognate response-element system was engineering using binding sites for the respective DBDs, inserted upstream a minimal TATA promoter to create an inducible gene expression system activated by SNIPRs utilizing humanized TFs. **(B)** Testing fully humanized SNIPRs. T cells expressing a SNIPR-BFP circuit were co-incubated with of either K562 or K562^CD19^ target cells for up to 72 hours and BFP output was measured using flow cytometry. **(C)** Testing receptor scaffold compatibility with humanized TFs. T cells engineered with mouse synNotch, human synNotch, or optimized hinge SNIPR circuits using humanized TFs were co-incubated with K562 or K562^CD19^ target cells for up to 72 hours and BFP output was measured by flow cytometry. **(D)** Target cell killing by a fully humanized SNIPR circuit. T cells expressing a SNIPR-CAR circuit were coincubated with K562 target cells for 72 hours. Target cells were cleared by 48 hours, as measured by DRAQ7 staining and flow cytometry. **(E)** In vivo assessment of SNIPR circuit function. NSG mice (5 per experimental group) were injected subcutaneously with 1×10^6^ K562^CD19/BCMA^ target cells into the left flank and 1×10^6^ K562^BCMA^ control cells into the right flank. 4 days post tumor injection, 6×10^6^ untransduced, BCMA CAR, or CD19 SNIPR-BCMA CAR circuit T cells (3×10^6^ each CD4+ and CD8+) were injected via tail vein, and tumor size was measured by caliper every few days. Statistics were calculated using one-way analysis of variance (ANOVA) with Dunnet’s test post hoc comparing untransduced T cells to BCMA CAR (**E** top, *) and Circuit T cells (**E** top, **) on Day 21, and comparing BCMA CAR T to circuit T cells (**E** bottom). *P ≤ 0.05, **P ≤ 0.01.

### *In vivo* testing of SNIPR-CAR circuits

Current challenges in CAR immunotherapy include the difficulty in defining a tumor with a single antigen. Systemic and unintended toxicity through on-target, off-tumor CAR activity has limited the clinical development of CARs and potent cytokine therapies (Ellis et al., 2021; Morgan et al., 2010). A multi-antigen recognition platform where a SNIPR binds a primary tumor antigen and drives expression of a CAR to a secondary antigen helps to mitigate risk of toxicity through more precise tumor recognition, and our humanized SNIPRs reduce the chance for immune rejection (Roybal et al., 2016b).

Next-generation humanized SNIPR-CAR circuits performed with high-fidelity during *in vitro* testing, but the question remained of their performance *in vivo*, where they would be exposed to a more diverse set of proteases and other environmental factors. To assess the performance and specificity of the optimized CD8α hinge SNIPRs *in vivo*, we examined the ability of SNIPR circuit T cells to control tumor growth in a dual-antigen xenograft model. Four days after implantation of CD19+/BCMA+ K562 tumors in the left flank and of BCMA+ K562 tumors in the right flank, NOD.Cg-*Prkdc^scid^II2rg^tm1Wjl^*/SzJ (NSG) mice were treated with untransduced T-cells, anti-BCMA CAR T cells, or anti-CD19 SNIPR circuit T cells containing either a Gal4-VP64- or HNF1A-p65-driven anti-BCMA CAR payload (**Fig. S5B, S5C**). The anti-BCMA CAR and anti-CD19 SNIPR - anti-BCMA CAR circuit T-cells controlled tumor growth in the left, dual-positive tumor, but only the BCMA CAR controlled tumor growth in the right, single positive tumor (**Fig. 5E, S5D**). A repeat experiment with the Gal4-VP64-driven circuit also revealed an absence of T cells in the right tumor and preferential SNIPR activation in the left tumor (**Fig. S5E-S5J**). These data support the potency and specificity of SNIPR circuits in an *in vivo* setting and represent the first successful demonstration of a humanized synthetic receptor circuit *in vivo*.

## DISCUSSION

From our investigations into the ECDs, TMDs, and JMDs of RIP receptors, we have constructed a large set of receptors that function like Notch and have begun to define the guidelines for the synthetic assembly of these receptors we call SNIPRs (**Fig. 6, Supplemental Tables S2, S3**). Overactive, inactive, and suboptimal core domains that control RIP all significantly reduce SNIPR performance, even when assembled with functional domains at other positions, suggesting that all three domains must be optimized for maximum ligand-dependent cleavage. We find that ECD specificity can be optimized by avoiding exposed protease sites and minimizing length, although SNIPR activity in response to alternative stimuli, such as T cell activation, may require direct observation to discover. We also find that additional TMDs and JMDs can be used to construct SNIPRs, and that TMDs and JMDs can be tuned through point mutations to meet individual clinical requirements, such as improving specificity or regulating levels of a delivered therapeutic. In addition, we find that SNIPRs containing suboptimal modules, such as the human synNotch ECD, can be improved through either direct ECD engineering, such as deletion of the NRR (**Fig. 2A, 2B**), or increasing activity in another module, such as the JMD. All three core SNIPR components, along with the LBD and TF, can impact receptor expression. Our systematic exploration of SNIPR parts has allowed us to identify receptors that are well-expressed and activate with high fidelity, two key features for robust cell therapy manufacturing and persistent activity in patients.

**Figure 6.**
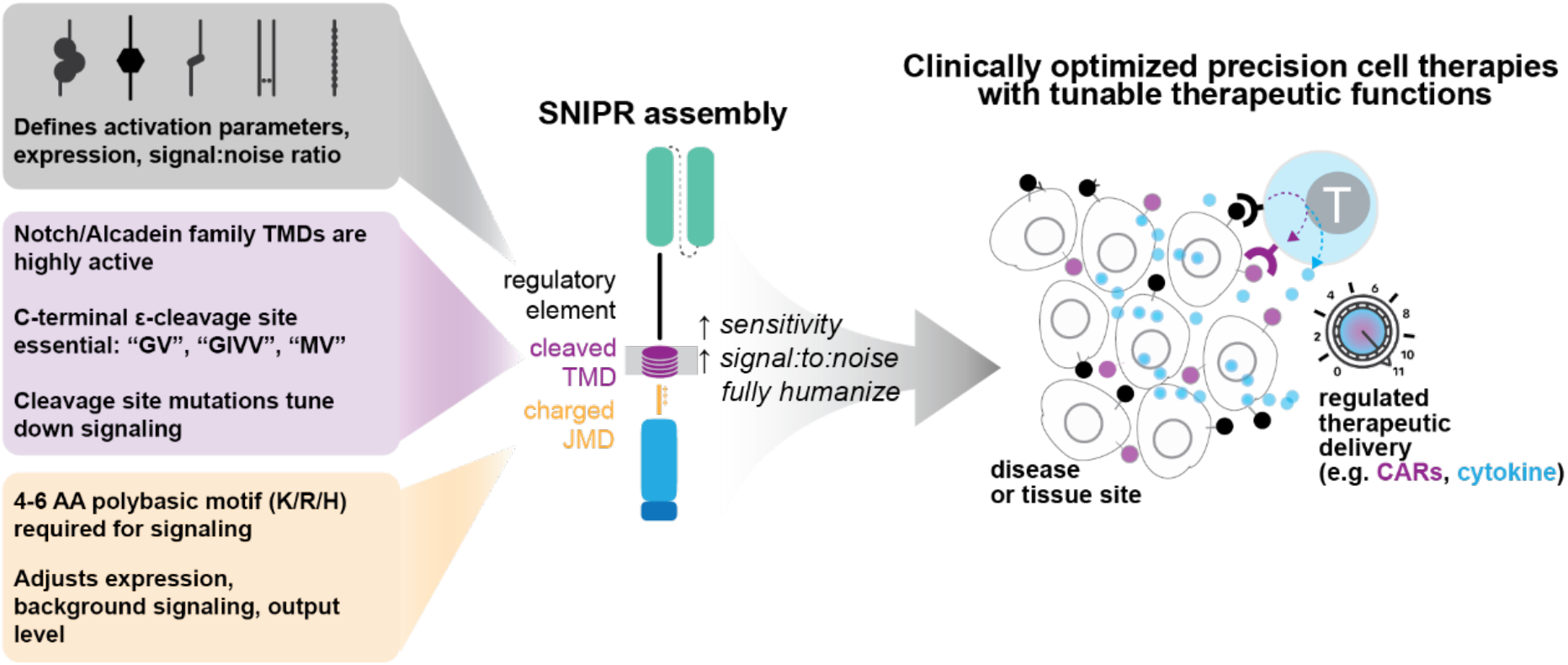
Design framework for next generation synthetic receptors for custom transcriptional regulation in therapeutic cells. Clinically relevant SNIPRs can be built through design of the receptor ECD, TMD, and JMD. The receptor ECD represents the first regulatory site and affects receptor activation parameters, expression, and stringency for ligand. Several known C-terminal motifs in the receptor TMD, commonly found in the Notch and Calsyntenin families, appear to be important for receptor signaling. Highly basic residues in the receptor JMD are required for signaling, and the choice of JMD can strongly affect receptor expression and output levels. By combining these elements, clinically relevant SNIPRs can be built that utilize fully human proteins and are compact, highly expressed, and regulatable. Our SNIPR design framework opens up the possibility to build truly customized precision cellular therapeutics.

We have found that many SNIPR ECDs that lack canonical regulatory domains such as the NRR remain functional. This result adds to previous screens of ECDs in a Notch context, which found that proteolytic switches with homology to Notch could substitute for the Notch NRR, albeit with a reduced signal-to-noise ratio (Hayward et al., 2019). In contrast, we find that several ECDs with no homology with Notch outperform it in the context of a synthetic receptor. One commonality between these functional ECDs is a relative lack of known protease cleavage sites. While a simple glycine-serine linker is a sufficient ECD, we find the addition of ADAM or MMP9 cleavage sites to this inherently unstructured linker leads to uncontrolled SNIPR activation. In addition, removal of known ADAM10 cleavage sites in the Robo1 and Notch1 ECDs improved the signal-to-noise ratios for SNIPRs utilizing these components. This discovery suggests that there is a large realm of permissive ECDs with a common mechanism of activation. Although most SNIPRs exclude known sites for ADAM protease cleavage, we find that all tested SNIPRs continue to rely on ADAM protease and γ-secretase activity (**Fig. S1E, S4B**). Our finding of enhanced SNIPR signaling during T cell activation may be explained by higher ADAM10 and ADAM17 activity (Li et al., 2007, Lambrecht et al., 2018), but the exact mechanism for either ligand-dependent or -independent activation for the diverse range of SNIPRs requires further study. Indeed, the mechanism of Notch activation through RIP is well-investigated, but the roles of other cellular processes, such as ubiquitination (Moretti et al., 2013), receptor endocytosis (Kandachar and Roegiers, 2012), and receptor trafficking (Yamamoto et al., 2009) remain unclarified and could also play a role in the activation of SNIPRs.

We were surprised to observe a lack of diversity in high-performing TMDs from our screen, having selected candidate TMDs from reported γ-secretase substrates (Haapasalo and Kovacs, 2011). This finding may be specific for SNIPRs expressed in human T cells, and SNIPRs containing non-functional TMDs from our screen may be more active when expressed in other tissue or cell types. Another possibility is that the cleavage efficiency of a particular TMD may have been evolutionarily selected for as a regulatory mechanism that favors prevention of spurious signaling rather than maximal activity.

From our systematic engineering of the SNIPR scaffold, we have built customizable receptor cores that provide not only the spatial discrimination afforded by previous synthetic receptors, such as synNotch, but are also more sensitive to lower antigen levels and more reliant on humanized components, thereby lowering the risk for immunogenicity. This added functionality is of clear benefit to current immunotherapies, such as CAR T cells, and should help provide a titrated therapeutic response while mitigating known issues of these technologies, such as premature T cell exhaustion and on-target/off-tumor systemic toxicity. For example, local titrated delivery of a potent cytokine, such as IL-12 (Lasek et al., 2014), to a tumor site using therapeutic cells may significantly improve efficacy and clinical outcomes as compared to the severe toxicity observed during systemic IV administration. These receptors should provide biomedical research with a comprehensive toolkit for directing a range of cell-based therapies to their intended targets combined with programmed localized therapeutic activity.

## EXPERIMENTAL PROCEDURES

### Receptor and Response Element Construct Design

Receptors were built by fusing the CD19 scFv (Porter et al., 2011), ALPPL2 M25^FYIA^ scFv (Hyrenius-Wittsten et al., 2021), HER2 4D5-8 scFv (Carter et al., 1992), EGFRviii 139 scFv (Morgan et al., 2012), or LaG17 nanobody (Fridy et al., 2014) to an extracellular domain comprised of: the human Notch1 (P46531) minimal regulatory region (Ile1427 to His1735), a truncated human Notch1 Notch Regulatory Region (Ile1427 to Glu1447, Thr1725 to His1735), a CD8α (P01732) hinge region (Thr138 to Asp182), a CD28 (P10747) hinge region (Ile114 to Pro152), a IgG4 hinge region, a OX40 hinge region, (the type III fibronectin domain from Robo1 (Q9Y6N7, Lys769 to Pro897), truncated CD8α hinges and fibronectin domains (as described), or Gly-Gly-Ser linkers of variable length (as described). All extracellular domains were fused to a transmembrane domain and intracellular juxtamembrane domain (as described), and a transcriptional element composed of Gal4 DBD VP64, Pax6(M1 to Ala139)-p65(Pro428 to Ser551), HNF1A(Met1 to Met283 with Thr-Cys-Arg linker)-p65(Asp361 to Ser551), or ZF-p65 (*14*). All receptors contain an N-terminal CD8α signal peptide (MALPVTALLLPLALLLHAARP) for membrane targeting and a myc-tag (EQKLISEEDL) for easy determination of surface expression with α-myc AF647 (Cell-Signaling #2233). The receptors were cloned into a modified pHR’SIN:CSW vector containing a PGK promoter for all primary T cell experiments. The pHR’SIN:CSW vector was also modified to make the response element plasmids. Five copies of the Gal4 DNA binding domain target sequence (GGAGCACTGTCCTCCGAACG), or four copies of the Pax6 consensus DBD recognition motif (ATTTTCACGCATGAGTGCACAG) and HNF1A DBD recognition motif (GTTAATNATTAAC) were cloned 5’ to a minimal synthetic pybTATA promoter. Also included in the response element plasmids is a PGK promoter that either constitutively drives expression of a fluorophore (mCitrine or mCherry) to easily identify transduced T cells or a SNIPR for single vector experimentation. Inducible CAR vectors contained CARs tagged N-terminally with FLAG-tag, and in some cases C-terminally with a t2a GFP system. All induced elements were cloned via a BamHI site in the multiple cloning site 3’ to the Gal4 response elements. All constructs were cloned via In-Fusion cloning (Takara # 638951).

### Primary Human T cell Isolation and Culture

Primary CD4+ and CD8+ T cells were isolated from anonymous donor blood after apheresis by negative selection (STEMCELL Technologies #15062 & 15063). Blood was obtained from Blood Centers of the Pacific (San Francisco, CA) as approved by the University Institutional Review Board. T cells were cryopreserved in RPMI-1640 (Thermo Fisher #11875093) with 20% human AB serum (Valley Biomedical Inc., #HP1022) and 10% DMSO. After thawing, T cells were cultured in human T cell medium consisting of X-VIVO 15 (Lonza #04-418Q), 5% Human AB serum and 10 mM neutralized N-acetyl L-Cysteine (Sigma-Aldrich #A9165) supplemented with 30 units/mL IL-2 (NCI BRB Preclinical Repository) for most experiments. For experiments involving the induction of Super IL-2, primary T-cells were maintained in human T cell media supplemented with IL-2 until experimentation, whereupon media was replaced with media without supplemented IL-2.

### Lentiviral Transduction of Human T cells

Pantropic VSV-G pseudotyped lentivirus was produced via transfection of Lenti-X 293T cells (Clontech #11131D) with a pHR’SIN:CSW transgene expression vector and the viral packaging plasmids pCMVdR8.91 and pMD2.G using Mirus Trans-IT Lenti (Mirus #MIR6606). Primary T cells were thawed the same day, and after 24 hours in culture, were stimulated with Human T-Activator CD3/CD28 Dynabeads (Life Technologies #11131D) at a 1:3 cell:bead ratio. At 48 hours, viral supernatant was harvested and the primary T cells were exposed to the virus for 24 hours. At day 5 post T cell stimulation, the Dynabeads were removed, and the T cells were sorted for assays with a Beckton Dickinson (BD) FACs ARIA II. Sorted T-cells were expanded until day 10 for *in vivo* assays and until day 14 for *in vitro* assays.

### Generation of Receptor Jurkat cells for Screening

E6-1 Jurkat T cells (ATCC# TIB-152) were lentivirally transduced with a reporter plasmid encoding a Gal4 driven tagBFP response element and a constitutively expressed mCitrine cassette. Reporter positive cells were sorted for mCitrine positivity and expanded. Individual cultures of reporter positive Jurkat T cells were lentivirally transduced in a 96 well plate with myc-tagged α-CD19 human SynNotch1 receptors with modified transmembrane or juxtamembrane domains. After viral transduction, the receptor transduction efficiency for each Jurkat cell population was measured with a BD FACSymphony Fortessa X-50 following staining with anti-myc AF647 (Cell-Signaling #2233).

### Cancer Cell Lines

The cancer cell lines used were K562 myelogenous leukemia cells (ATCC #CCL-243). K562s were lentivirally transduced to stably express either human CD19 at equivalent levels as Daudi tumors (ATCC #CCL-213), BCMA, or both BCMA and CD19. CD19 levels were determined by staining the cells with α-CD19 APC (Biolegend #302212). BCMA levels were determined by staining the cells with α-BCMA APC (Biolegend #357505). All cell lines were sorted for expression of the transgenes.

### MCAM BiTE Production

MCAM BiTE was produced from transfecting LentiX-293T cells with a pHR’SIN:CSW transgene expression vector. 293T media was replaced with T-cell media 24 hours after transfection. MCAM BiTE was harvested 48 hours post-media replacement by collecting supernatant and removing 293T cells via centrifugation.

### Doxycycline inducible ALPPL2

A clonal line of K562 cells expressing a doxycycline-inducible FLAG-tagged ALPPL2 cassette was treated with doxycycline (Abcam) at doses ranging between 0.1-100 ng/mL for 24 hours prior to co-incubation with T cells. Surface expression levels were assessed by flow cytometry through the FLAG-tag on ALPPL2 prior to assay.

### *In vitro* SNIPR Activation Assays

For all *in vitro* SNIPR activation assays, 1×10^5^ T cells or Jurkat T cells were co-cultured with target cells at a 1:1 ratio in 96 well round bottom plates (VWR). To exogenously activate T-cells, 100 ng/mL PMA (Sigma #P1575) or MCAM BiTEs were added to cocultures. When activating a co-expressed ALPPL2 CAR, ALPPL2+ K562 cells were added to the co-culture in a 1:1 ratio with T cells. The cultures were analyzed at the time points indicated for reporter activation using a BD FACSymphony Fortessa X-50. All flow cytometry analysis was performed in FlowJo software (BD). TMD sequence alignment was performed using ClustalX and visualized using Jalview.

### Super IL-2 Induction Assays

Primary CD4+ were stained with Cell Trace Violet (Thermo Fisher #C34557) and stimulated with irradiated K562 or CD19+ K562 target cells in human T cell media without IL-2 supplementation. Supernatant was harvested at the indicated timepoints and IL-2 levels in the supernatant were measured via IL-2 Human Instant ELISA kit (Thermo Fisher #BMS221INST). T-cell proliferation was also measured at the indicated timepoints using a BD FACSymphony Fortessa X-50.

### *In vivo* assays

NOD.Cg-*Prkdc^scid^II2rg^tm1WJl^*/SzJ (NSG) mice were implanted with either 1×10^6^ K562^CD19+/BCMA+^ tumor cells subcutaneously in the left flank alone or with an additional 1×10^6^ K562^BCMA+^ tumor cells subcutaneously in the right flank. Four days after tumor implantation, 2.5 or 3×10^6^ engineered primary human CD4+ and CD8+ T cells (total of 5 or 6×10^6^ T cells) were intravenously infused through tail vein injection. Tumor size was monitored via caliper regularly.

## SUPPLEMENTARY MATERIALS

Tables S1-S3

Figs. S1-S5

## ACKNOWLEDGEMENTS

I.Z. is funded by an F30 fellowship from the National Cancer Institute (5F30CA250247-02). R.L. was supported by a T32 training grant (5T32AI07334-29). K.T.R. is funded by the Parker Institute for Cancer Immunotherapy, the UCSF Helen Diller Family Comprehensive Cancer Center, the Chan Zuckerberg Biohub, an NIH Director’s New Innovator Award (DP2 CA239143), Cancer Research UK, and the Kleberg Foundation. This work was supported by NIH grant R01EB029483 (K.T.R., A.S.K.). A.S.K. acknowledges funding from a DARPA Young Faculty Award (D16AP00142), NIH Director’s New Innovator Award (1DP2AI131083), and DoD Vannevar Bush Faculty Fellowship (N00014-20-1-2825). We acknowledge the PFCC supported in part by Grant NIH P30 DK063720 and by the NIH S10 Instrumentation Grant S10 1S10OD021822-01.

## AUTHOR CONTRIBUTIONS

I.Z. designed the study, designed and performed experiments and vector construction, analyzed data, and wrote the manuscript; R.L. conceived and designed the study, designed and performed experiments and vector construction, analyzed data, and wrote the manuscript. A.H-W. performed and analyzed *in vivo* experiments; D.P. and D.V.I. designed vectors and performed experiments related to synthetic zinc fingers; J.A. performed *in vitro* experiments. A.S.K. conceived and designed the study regarding synthetic zinc finger transcription factors. K.T.R. conceived and designed the study, designed experiments, and wrote the manuscript.

