## Supplemental Material for "Design and modular assembly of synthetic intramembrane proteolysis receptors for custom gene regulation in therapeutic cells"

**SUPPLEMENTAL DATA**

**Supplemental Table S1.** Sequences of screened TMDs and JMDs. Sequences of screened TMDs and JMDs. TMDs and JMDs were sourced from known γ-secretase substrates (*19*), and Notch1 homologs. TMD sequences were extracted from Uniprot annotations and were extended to the first basic residue (R, K, or H). Two TMDs from proteins not believed to be γ-secretase substrates were selected for negative control. JMD sequences began immediately c-terminal to the TMD and ended immediately prior to 3 non-basic residues.

| Gene | ID | TMD | JMD |
|---|---|---|---|
| CLSTN1 | O94985 | ATVVIVVCVSFLVFMIILGVF | RIRAAHRRTMR |
| CLSTN2 | Q9H4D0 | IATVVIIISVCMLVFVVAMGVY | RVRIAHQH |
| APLP1 | P51693 | AVSGLLIMGAGGGSLIVLSMLLL | RRKK |
| APLP2 | Q06481 | SALIGLLVIAVAIATVIVISLVML | RKR |
| LRP8 | Q14114 | VIGIIVPIVVIALLCMSGYLIW | RNWKRKNTK |
| APP | P05067 | GAIIGLMVGGVVIATVIVITLVML | KKK |
| BTC | P35070 | ILVICLIAVMVVFIILVIGVCTCC | HPLRKRRKRKK K |
| TGBR3 | Q03167 | MGIAFAAFVIGALLTGALWYIYS | H |
| SPN | P16150 | GMLPVAVLVALLAVIVLVALLLLW | RRRQKRR |
| CD44 | P16070 | WLIILASLLALALILAVCIAVNS | RRRCGQKKK |
| CSF1R | P07333 | VVVACMSIMALLLLLLLLLY | KYKQKPK |
| CXCL16 | Q9H2A7 | VPVLCLLAIIFILTAALSYVLC | KRRR |
| CX3CL1 | P78423 | AVGLLAFLGLLFCLGVAMFTYQSLQ GCP | RK |
| DCC | P43146 | LLVIIVVTVGVITVLVVVIVAVICT | RR |
| DLL1 | O00548 | VAVCAGVILVLMLLLGCAAVVVCV | RLRLQKHR |
| DSG2 | Q14126 | LGPAAIALMILAFLLLLLVPLLLLMC | HCGKGAK |
| DNER | Q8NFT8 | IIIGALCVAFILMLIILIVGIC | RISR |
| DAG1 | Q14118 | YLHTVIPAVVVAAILLIAGIIAMICY | RKKRKGK |
| CDH1 | P12830 | ILGILGGILALLILILLLLLFL | RRR |
| EPCAM | P16422 | AGVIAVIVVVVIAVVAGIVVLVIS | RKKRMAKYEK |
| EPHA4 | P54764 | VLLVSVSGSVVLVVILIAAFVIS | RRRSKYSKAK |
| EPHB2 | P29323 | IIGSSAAGLVFLIAVVVIAIVCN | RR |
| EFNB1 | P98172 | VALFAAVGAGCVIFLLIIIFLTVLLL | KLRKRHRKH |
| EFNB2 | P52799 | GIASGCIIFIVIIITLVVLLL | KYRRRHRKH |
| ERBB4 | Q15303 | LIAAGVIGGLFILVIVGLTFAVYV | RRKSIKKKRALR R |
| GHR | P10912 | FPWLLIIIFGIFGLTVMLFVFLFS | KQQRIK |
| HLA-A | P01892 | VGIIAGLVLFGAVITGAVVAAVMW | RRK |
| IFNAR2 | P48551 | IGGIITVFLIALVLTSTIVTL | K |

| | | | |
|---|---|---|---|
| IGF1R | P08069 | LIIALPVAVLLIVGGLVIMLYVFH | RKR |
| IL1R1 | P14778 | HMIGICVTLTVIIVCSVFIY | KIFK |
| IL1R2 | P27930 | ASSTFSWGIVLAPLSLAFLVLGGIWM | HRRCKHRTGK |
| IL6R | P08887 | TFLVAGGSLAFGTLLCIAIVL | RFKKTWKLRAL KEGK |
| INSR | P06213 | IIIGPLIFVFLFSVVIGSIYLFL | RKR |
| ERN1 | O75460 | MATIILSTFLLIGWVAFIITYPLSM | H |
| ERN2 | Q76MJ5 | QDLLAASLTAVLLGGWILFVM | R |
| JAG2 | Q9Y219 | GLLVPVLCGAFSVLWLACVVLCVWW T | RKRRKERERSR LPR |
| KCNE1 | P15382 | ALYVLMVLGFFGFFTLGIMLSYI | RSKKLEH |
| KCNE2 | Q9Y6J6 | VILYLMVMIGMFSFIIVAILVSTV | KSKRREH |
| KCNE3 | Q9Y6H6 | YMYILFVMFLFAVTVGSLILGYT | RSRKVDKR |
| KCNE4 | Q8WWG 9 | YFYILVVMSFYGIFLIGIMLGYM | KSKRREKK |
| KL | Q9UEF7 | LLAFIAFLFFASIISLSLIFYYS | KKGRRSYK |
| CHL1 | O00533 | FIGLMCAIALLTLLLLTVCFV | KRNRGGK |
| PTPRF | P10586 | VTGPVLAVILIILIVIAILLF | KRKRTH |
| LRP1 | Q07954 | HIASILIPLLLLLLLVLVAGVVFWY | KRR |
| LRP1B | Q9NZR2 | AIIVPLVLLVTLITTLVIGLVLC | KRKRRTKTIRR |
| LRP2 | P98164 | AVAVLLTILLIVVIGALAIAGFF | HYRR |
| LRP6 | O75581 | TNTVGSVIGVIVTIFVSGTVYFICQ | R |
| MUC1 | P15941 | WGIALLVLVCVLVALAIVYLIALAVCQ C | RRK |
| CDH2 | P19022 | IIAILLCIIILLILVLMFVVWM | KRRDKERQAK |
| SCN1B | Q07699 | IMMYVLIVVLTIWLVAEMIYCY | KK |
| SCN2B | O60939 | VIVGASVGGFLAVVILVLMVV | KCVRRKKEQK |
| SCN3B | Q9NY72 | IMMYILLVFLTLWLLIEMIYCY | RKVSK |
| SCN4B | Q8IWT1 | TLIILAVVGGVIGLLILILLI | KK |
| NECTIN1 | Q15223 | IIGGVAGSILLVLIVVGGIVVAL | RRRRHTFK |
| NRG1 | Q02297 | VLTITGICIALLVVGIMCVVAYC | KTKKQRKKLHD RLR |
| NRG2 | O14511 | VLTITGICVALLVVGIVCVVAYC | KTKKQRKQMHN HLR |
| NOTCH1 | P46531 | FMYVAAAAFVLLFFVGCGVLLS | RKRRR |
| NOTCH2 | Q04721 | LLYLLAVAVVIILFIILLGVIMA | KRKRKH |
| NOTCH3 | Q9UM47 | LPLLVAGAVLLLVILVLGVMVA | RRKREH |
| NOTCH4 | Q99466 | PVLCSPVAGVILLALGALLVLQLI | RRRRREH |
| NPR3 | P17342 | SAVTGIVVGALLGAGLLMAFYFF | RKKYR |
| Nradd | Q8CJ26 | IIPVYCALLATVILGLLAYVAF | KCWRSHKQR |
| NGFR | P08138 | LIPVYCSILAAVVVGLVAYIAF | KR |
| PAM | P19021 | VPVVLITTLLVIPVVVLLAIAIFI | RWKKSR |
| PLXDC2 | Q6UX71 | GLIIGILILVLIVATAILVTVYMY | HH |

| | | | |
|---|---|---|---|
| PKHD1 | P08F94 | IILAASLSSVASWLALSCLVCCWL | KRSKSRKTK |
| PCDHA4 | Q9UN74 | VYLIIAICAVSSLLVLTLLLYTAL | R |
| PCDHGC3 | Q9UN70 | LLLSLILVSVGFVVTVFGVIIF | KVYKWKQSR |
| PTPRZ1 | P23471 | AVIPLVIVSALTFICLVVLVGILIYW | RK |
| AGER | Q15109 | LALGILGGLGTAALLIGVILWQ | RRQRR |
| PTPRK | Q15262 | IAGISAGILVFILLLLVVILIV | KKSKLAKKRK |
| PTPRM | P28827 | IAGVIAGILLFVIIFLGVVLVM | KKRKLAKKRK |
| ROBO1 | Q9Y6N7 | AFIAGIGAACWIILMVFSIWLY | RHRKKR |
| SORCS3 | Q9UPU3 | AMLMLLSVVFVGLAVFLIYKF | KRK |
| SORCS1 | Q8WY21 | GSAMLMLLSVVFVGLAVFVIY | KFKRR |
| SORL1 | Q92673 | AVVVPILFLILLSLGVGFAILYT | KHRR |
| SORT1 | Q99523 | SVPIILAIVGLMLVTVVAGVLIV | KK |
| SDC1 | P18827 | GVIAGGLVGLIFAVCLVGFMLY | RMKKK |
| SDC2 | P34741 | VLAAVIAGGVIGFLFAIFLILLLVY | RMRKK |
| SDC3 | O75056 | AVIVGGVVGALFAAFLVTLLIY | RMKKK |
| TIE1 | P35590 | QLILAVVGSVSATCLTILAALLTLVCI | RR |
| TYR | P14679 | WLLGAAMVGAVLTALLAGLVSLLC | RHKRK |
| TYRP1 | P17643 | IIAIAVVGALLLVALIFGTASYLI | RARR |
| DCT | P40126 | LLVVMGTLVALVGLFVLLAFLQY | RRLRK |
| VASN | Q6EMK4 | LLIAPALAAVLLAALAAVGAAYCV | RRGR |
| CDH5 | P33151 | AVVAILLCILTITVITLLIFL | RRRLRKQARAHGK |
| FLT1 | P17948 | LITLTCTCVAATLFWLLLTLFI | RKMKR |
| VLDLR | P98155 | AAWAILPLLLLVMAAVGGYLMW | R |
| NOTCH1_D. rerio | P46530 | MYPMFLVLLALAVLALAAVGVVVS | Not Screened |
| NOTCH1_D. melanogaster | P07207 | VITGIILVIIALAFFGMVLSTQ | Not Screened |
| NOTCH1_X. laevis | P21783 | PMLSMLVIPLLIIFVFMMVIVN | Not Screened |
| NOTCH1_G. gallus | F1NZ70 | PMYVVVAALVLLAFIGVGVLVS | Not Screened |
| NCSTN (Negative control) | Q92542 | LITLTVGFGILIFSLIVTYCINA | Not Screened |
| CD147 (Negative control) | P35613 | ALWPFLGIVAEVLVLVTIIFIYE | Not Screened |

**Supplemental Table S2.** Components of highlighted SNIPRs.

| Component ID | Description | Sequence |
|---|---|---|
| CD19 scFv | CD8αsignalpeptide_myc-tag_CD19scFv | MALPVTALLLPLALLLHAARPEQKLISEEDLDI QMTQTTSSLSASLGDRVTISCRASQDISKYL NWYQQKPDGTVKLLIYHTSRLHSGVPSRFS GSGSGTDYSLTISNLEQEDIATYFCQQGNTL PYTFGGGTKLEITGGGGSGGGGSGGGGSE VKLQESGPGLVAPSQSLSVTCTVSGVSLPD YGVSWIRQPPRKGLEWLGVIWGSETTYYN SALKSRLTIIKDNSKSQVFLKMNSLQTDDTA IYYCAKHYYYGGSYAMDYWGQGTSVTVSS |
| ALPPL2 scFv | CD8αsignalpeptide_myc-tag_ALPPL2scFv | MALPVTALLLPLALLLHAARPEQKLISEEDL QVQLQQSGGGLVKPGGSLRLSCAASGFTF SSYAMHWVRQAPGKGLEWVAVISYDGSN KYYADSVKGRFTISRDNSKNTLYLQMDSLR AEDTAVYYCAKEGDSSRWSYDLWGRGTLV TVSSGGGGSGGGGSGGGGSQSALTQPAS VSGSPGQSITISCTGTSSDVGGYNYVSWYQ QHPGKAPKVMIYDVTNRPSGVSNRFSGSKS GNTASLTISGLQAEDEADYYCS SYTIASTLVVFGGGTKLTVL |
| LaG17 nanobody | CD8αsignalpeptide_myc-tag_LaG17nanbody | MALPVTALLLPLALLLHAARPEQKLISEEDLM ADVQLVESGGGLVQAGGSLRLSCAASGRTI SMAAMSWFRQAPGKEREFVAGISRSAGSA VHADSVKGRFTISRDNTKNTLYLQMNSLKAE DTAVYYCAVRTSGFFGSIPRTGTAFDYWGQ GTQVTVS |
| EGFRviii scFv (139) | CD8αsignalpeptide_myc-tag_EGFRviiiscFv | MALPVTALLLPLALLLHAARPEQKLISEEDLDI Q MTQSPSSLSASVGDRVTITCRASQGIRNNLA W YQQKPGKAPKRLIYAASNLQSGVPSRFTGSG S GTEFTLIVSSLQPEDFATYYCLQHHSYPLTSG G GTKVEIKGSTSGSGKPGSGEGSEVQVLESG G GLVQPGGSLRLSCAASGFTFSSYAMSWVRQ A PGKGLEWVSAISGSGGSTNYADSVKGRFTIS R |

| | | |
|---|---|---|
| | | DNSKNTLYLQMNSLRAEDTAVYYCAGSSGW<br>S<br>EYWGQGTLVTVSS |
| HER2 scFv (4D5-8) | CD8αsignalpeptide_myc-tag_HER2scFv | MALPVTALLLPLALLLHAARPEQKLISEEDLDI<br>Q<br>MTQSPSSLSASVGDRVTITCRASQDVNTAVA<br>W<br>YQQKPGKAPKLLIYSASFLYSGVPSRFSGSR<br>S<br>GTDFTLTISSLQPEDFATYYCQQHYTTPPTFG<br>QGTKVEIKRTGSTSGSGKPGSGEGSEVQLVE<br>SGGGLVQPGGSLRLSCAASGFNIKDTYIHWV<br>RQAPGKGLEWVARIYPTNGYTRYADSVKGR<br>F<br>TISADTSKNTAYLQMNSLRAEDTAVYYCSRW<br>GGDGFYAMDVWGQGTLVTVSSGS |
| mN1 ECD | Notch1 NRR from murine Notch1 | ILDYSFTGGAGRDIPPPQIEEACELPECQVD<br>AGNKVCNLQCNNHACGWDGGDCSLNFND<br>PWKNCTQSLQCWKYFSDGHCDSQCNSAG<br>CLFDGFDCQLTEGQCNPLYDQYCKDHFSD<br>GHCDQGCNSAECEWDGLDCAEHVPERLA<br>AGTLVLVVLLPPDQLRNNSFHFLRELSHVLH<br>TNVVFKRDAQGQQMIFPYYGHEEELRKHPI<br>KRSTVGWATSSLLPGTSGGRQRRELDPMD<br>IRGSIVYLEIDNRQCVQSSSQCFQSATDVA<br>AFLGALASLGSLNIPYKIEAVKSEPVEPPLPSQ |
| hN1 ECD | Notch1 NRR from human Notch1 | ILDYSFGGGAGRDIPPPLIEEACELPECQEDA<br>GNKVCSLQCNNHACGWDGGDCSLNFNDPW<br>KNCTQSLQCWKYFSDGHCDSQCNSAGCLFD<br>GFDCQRAEGQCNPLYDQYCKDHFSDGHCD<br>QGCNSAECEWDGLDCAEHVPERLAAGTLVV<br>VVLMPPEQLRNSSFHFLRELSRVLHTNVVFK<br>RDAHGQQMIFPYYGREEELRKHPIKRAAEG<br>WAAPDALLGQVKASLLPGGSEGGRRRRELD<br>PMDVRGSIVYLEIDNRQCVQASSQCFQSATD<br>VAAFLGALASLGSLNIPYKIEAVQSETVEPPP<br>PAQLH |
| hN1 ΔNRR ECD | hN1 ECD with NRR deleted and flanking regions fused | ILDYSFGGGAGRDIPPPLIEETVEPPPPAQLH |
| hN2 ECD | Notch2 NRR from human Notch2 | LYTAPPSTPPATCLSQYCADKARDGVCDEAC<br>NSHACQWDGGDCSLTMENPWANCSSPLPC |

| | | |
|---|---|---|
| | | WDYINNQCDELCNTVECLFDNFECQGNSKT CKYDKYCADHFKDNHCDQGCNSEECGWDG LDCAADQPENLAEGTLVIVVLMPPEQLLQDA RSFLRALGTLLHTNLRIKRDSQGELMVYPYY GEKSAAMKKQRMTRRSLPGEQEQEVAGSK VFLEIDNRQCVQDSDHCFKNTDAAAALLAS HAIQGTLSYPLVSVVSESLTPERTQ |
| hN2 ΔNRR ECD | hN2 ECD with NRR deleted and flanking regions fused | LYTAPPSTPPATSLTPERTQ |
| hN3 ECD | Notch3 NRR from human Notch3 | CPRAACQAKRGDQRCDRECNSPGCGWDG GDCSLSVGDPWRQCEALQCWRLFNNSRCD PACSSPACLYDNFDCHAGGRERTCNPVYEK YCADHFADGRCDQGCNTEECGWDGLDCAS EVPALLARGVLVLTVLLPPEELLRSSADFLQR LSAILRTSLRFRLDAHGQAMVFPYHRPSPGS EPRARRELAPEVIGSVVMLEIDNRLCLQSPEN DHCFPDAQSAADYLGALSAVERLDFPYPLRD VRGE |
| hN3 ΔNRR ECD | hN3 ECD with NRR deleted and flanking regions fused | APAAAPEVSEEPRPLEPPEPSVPL |
| hN4 ECD | Notch4 NRR from human Notch4 | KPGAKGCEGRSGDGACDAGCSGPGGNWD GGDCSLGVPDPWKGCPSHSRCWLLFRDGQ CHPQCDSEECLFDGYDCETPPACTPAYDQY CHDHFHNGHCEKGCNTAECGWDGGDCRP EDGDPEWGPSLALLVVLSPPALDQQLFALAR VLSLTLRVGLWVRKDRDGRDMVYPYPGARA EEKLGGTRDPTYQERAAPQTQPLGKETDSL SAGFVVVMGVDLSRCGPDHPASRCPWDPG LLLRFLAAMAAVGALEPLLPGPLLAVHPHAG TAPPANQLPW |
| hN4 ΔNRR ECD | hN4 ECD with NRR deleted | VHPHAGTAPPANQLPW |
| Robo1 ECD | Fn-III domain and n-terminal JMD from human Robo1 | KTLEEAPSAPPQGVTVSKNDGNGTAILVSWQ PPPEDTQNGMVQEYKVWCLGNETRYHINKT VDGSTFSVVIPFLVPGIRYSVEVAASTGAGSG VKSEPQFIQLDAHGNPVSPEDQVSLAQQISD VVKQP |
| Fn-III ECD | Fn-III domain from human Robo1 | KTLEEAPSAPPQGVTVSKNDGNGTAILVSW QPPPEDTQNGMVQEYKVWCLGNETRYHINK TVDGSTFSVVIPFLVPGIRYSVEVAASTGAGS GVKSEPQFIQLDVVKQP |

| | | |
|---|---|---|
| 3X FLAG ECD | 3xFLAG-tag tether | DYKDHDGDYKDHDIDYKDDDDKEPPPPAQLH |
| CD8α ECD | CD8α hinge ECD | TTTPAPRPPTPAPTIASQPLSLRPEACRPAAGGAVHTRGLDFACD |
| CD8α ECD, variant 1 | truncated CD8α hinge ECD, version 1 | TTTPAPRPPTPAPTIASQPLSLRPEA |
| CD8α ECD, variant 2 | Optimized CD8α hinge ECD | TTTPAPRPPTPAPTIASQPLSLRPEAC |
| CD8α ECD, variant 3 | truncated CD8α hinge ECD, version 3 | RPAAGGAVHTRGLDFACD |
| CD8α ECD, variant 4 | truncated CD8α hinge ECD, version 4 | CRPAAGGAVHTRGLDFACD |
| CD28 ECD | CD28 hinge ECD | IEVMYPPPYLDNEKSNGTIIHVKGKHLCPSPLFPGPSKP |
| IgG4 ECD | (G4S)3_IgG4 hinge ECD | GGGGSGGGGSGGGGSESKYGPPCPPCP |
| OX40 ECD | OX40 hinge ECD | LHCVGDTYPSNDRCCHECRPGNGMVSRCSRSQNTVCRPCGPGFYNDVVSSKPCKPCTWCNLRSGSERKQLCTATQDTVCRCRAGTQPLDSYKPGVDCAPCPPGHFSPGDNQACKPWTNCTLAGKHTLQPASNSSDAICEDRDPPATQPQETQGPPARPITVQPTEAWPRTSQGPSTRPVEVPGGRA |
| mN1 TMD | murine Notch1 TMD | LMYVAAAAFVLLFFVGCGVLLS |
| hN1 TMD | human Notch1 TMD | FMYVAAAAFVLLFFVGCGVLLS |
| hN1 TMD_G318A | human Notch1 TMD with c-terminal G-->A mutant | FMYVAAAAFVLLFFVGCAVLLS |
| hN1 TMD_V319A | human Notch1 TMD with c-terminal V-->A mutant | FMYVAAAAFVLLFFVGCGALLS |
| hN2 TMD | human Notch2 TMD | LLYLLAVAVVIILFIILLGVIMA |
| hN3 TMD | human Notch3 TMD | LPLLVAGAVLLLVILVLGVMVA |
| hN4 TMD | human Notch4 TMD | PVLCSPVAGVILLALGALLVLQLI |
| Robo1 TMD | Robo1 TMD | AFIAGIGAACWIILMVFSIWLY |
| Robo1 TMD_GV | Robo1 TMD with "IW" mutated to "GV" | AFIAGIGAACWIILMVFSGVLY |
| CLSTN2 TMD | CLSTN2 TMD | IATVVIIISVCMLVFVVAMGVY |

| | | |
|---|---|---|
| AGER TMD | AGER TMD | LALGILGGLGTAALLIGVILWQ |
| AGER TMD_LVS | AGER TMD with "ILWQ" mutated to "LVS" | LALGILGGLGTAALLIGVLVS |
| Robo1 JMD | Robo1 c-terminal JMD | RHRKKR |
| N1 JMD | Notch1 c-terminal JMD | RKRRR |
| N2 JMD | Notch2 c-terminal JMD | KRKRKH |
| N3 JMD | Notch3 c-terminal JMD | RKRREH |
| N4 JMD | Notch4 c-terminal JMD | RRRRREH |
| CLSTN2 JMD | CLSTN2 JMD | RVRIAHQH |
| AGER JMD | AGER JMD | RRQRR |
| PTPRF JMD | PTPRF JMD | KRKRTH |
| NRG1 JMD | NRG1 JMD | KTKKQRKKLHDRLR |
| G4VP64 | Gal4DBD_VP64 | MKLLSSIEQACDICRLKKLKCSKEKPKCAK CLKNNWECRYSPKTKRSPLTRAHLTEVES RLERLEQLFLLIFPREDLDMILKMDSLQDIK ALLTGLFVQDNVNKDAVTDRLASVETDMP LTLRQHRISATSSSEESSNKGQRQLTVSA AAGGSGGSGGSDALDDFDLDMLGSDALD DFDLDMLGSDALDDFDLDMLGSDALDDFD LDMLGS |
| Pax6 | Pax6 DBD | MQNSHSGVNQLGGVFVNGRPLPDSTRQK IVELAHSGARPCDISRILQVSNGCVSKILGR YYETGSIRPRAIGGSKPRVATPEVVSKIAQ YKRECPSIFAWEIRDRLLSEGVCTNDNIPS VSSINRVLRNLASEKQQMGA |
| HNF1A | HNF1A DBD | MVSKLSQLQTELLAALLESGLSKEALLQAL GEPGPYLLAGEGPLDKGESCGGGRGELA ELPNGLGETRGSEDETDDDGEDFTPPILK ELENLSPEEAAHQKAVVETLLQEDPWRVA KMVKSYLQQHNIPQREVVDTTGLNQSHLS QHLNKGTPMKTQKRAALYTWYVRKQREV AQQFTHAGQGGLIEEPTGDELPTKKGRRN RFKWGPASQQILFQAYERQKNPSKEERET LVEECNRAECIQRGVSPSQAQGLGSNLVT EVRVYNWFANRRKEEAFRHKLAM |

| | | |
|---|---|---|
| p65(361-551) | NFkB p65 transactivation domain | DEFPTMVFPSGQISQASALAPAPPQVLPQAPAPAPAPAMVSALAQAPAPVPVLAPGPPQAVAPPAPKPTQAGEGTLSEALLQLQFDDEDLGALLGNSTDPAVFTDLASVDNSEFQQLLNQGIPVAPHTTEPMLMEYPEAITRLVTGAQRPPDPAPAPLGAPGLPNGLLSGDEDFSSIADMDFSALLSQISS |
| p65(428-551) | NFkB p65 transactivation domain truncation | PTQAGEGTLSEALLQLQFDDEDLGALLGNSTDPAVFTDLASVDNSEFQQLLNQGIPVAPHTTEPMLMEYPEAITRLVTGAQRPPDPAPAPLGAPGLPNGLLSGDEDFSSIADMDFSALLSQISS |
| ZF6 | synthetic zinc finger DNA binding domain (synTF) 6 | (Israni et al., 2021) |
| ZF10 | synthetic zinc finger DNA binding domain (synTF) 10 | (Israni et al., 2021) |

**Supplemental Table S3.** Highlighted SNIPR designs.

| Figure # | SNIPR ID | Receptor | LBD | ECD | TMD | JMD | TF |
|---|---|---|---|---|---|---|---|
| S1B | synNotch | murine SynNotch | CD19 scFv | mN1 ECD | mN1 TMD | N1 JMD | G4VP64 |
| S1B, S1C, 2A-B, S3D | hsNotch1 LNR | human SynNotch1 | CD19 scFv | hN1 ECD | hN1 TMD | N1 JMD | G4VP64 |
| S1B | hsNotch2 LNR | human SynNotch2 | CD19 scFv | hN2 ECD | hN2 TMD | N2 JMD | G4VP64 |
| S1B | hsNotch3 LNR | human SynNotch3 | CD19 scFv | hN3 ECD | hN3 TMD | N3 JMD | G4VP64 |
| S1B | hsNotch4 LNR | human SynNotch4 | CD19 scFv | hN4 ECD | hN4 TMD | N4 JMD | G4VP64 |
| 1B, S1D, S3C | synRobo | synthetic Robo1 receptor | CD19 scFv | Robo1 ECD | Robo1 TMD | Robo1 JMD | G4VP64 |
| 1B, S1D | synRobo/Notch | synthetic Robo1 with Notch1 TMD/JMD | CD19 scFv | Robo1 ECD | hN1 TMD | N1 JMD | G4VP64 |
| 1B, S1D-E | FnIII-Notch | Robo1 Fn-III domain ECD with Notch1 TMD/JMD | CD19 scFv | Fn-III ECD | hN1 TMD | N1 JMD | G4VP64 |
| 1C | (GGS)3_SNIPR | 9AA GGS linker with Notch1 TMD/JMD | CD19 scFv | (GGS)3 | hN1 TMD | N1 JMD | G4VP64 |
| 1C | (GGS)6_SNIPR | 18AA GGS linker with Notch1 TMD/JMD | CD19 scFv | (GGS)6 | hN1 TMD | N1 JMD | G4VP64 |
| 1C, S1E | (GGS)9_SNIPR | 27AA GGS linker with Notch1 TMD/JMD | CD19 scFv | (GGS)9 | hN1 TMD | N1 JMD | G4VP64 |
| 1C | (GGS)12_SNIPR | 36AA GGS linker with Notch1 TMD/JMD | CD19 scFv | (GGS)12 | hN1 TMD | N1 JMD | G4VP64 |
| 1C | (GGS)15_SNIPR | 45AA GGS linker with | CD19 scFv | (GGS)15 | hN1 TMD | N1 JMD | G4VP64 |

| | | | | | | | |
|---|---|---|---|---|---|---|---|
| | | Notch1 TMD/JMD | | | | | |
| 1C | (GGS)18_ SNIPR | 54AA GGS linker with Notch1 TMD/JMD | CD19 scFv | (GGS)18 | hN1 TMD | N1 JMD | G4VP 64 |
| 2A | ADAM17 site | exposed ADAM17 site in ECD | CD19 scFv | (GGS)3(SPLAQAVRSSSR)(GGS)3 | hN1 TMD | N1 JMD | G4VP 64 |
| 2A, S2A | FAP site | exposed FAP site in ECD | --------------- | (GGGGSASGPAGPAGGGSGGSA)4 | hN1 TMD | N1 JMD | G4VP 64 |
| S2A | FAP site with scFv | exposed FAP site in ECD with scFv attached | CD19 scFv | (GGGGSASGPAGPAGGGSGGSA)4 | hN1 TMD | N1 JMD | G4VP 64 |
| 2A | 3xFLAG-tag tether | 3xFLAG-tag in ECD | CD19 scFv | 3X FLAG ECD | hN1 TMD | N1 JMD | G4VP 64 |
| 2A-B, S2F | Notch1 ΔNRR | human SynNotch1 with NRR deleted | CD19 scFv | hN1 ΔNRR ECD | hN1 TMD | N1 JMD | G4VP 64 |
| S2F | Notch2 ΔNRR | human SynNotch2 with NRR deleted | CD19 scFv | hN2 ΔNRR ECD | hN2 TMD | N2 JMD | G4VP 64 |
| S2F | Notch3 ΔNRR | human SynNotch3 with NRR deleted | CD19 scFv | hN3 ΔNRR ECD | hN3 TMD | N3 JMD | G4VP 64 |
| S2F | Notch4 ΔNRR | human SynNotch4 with NRR deleted | CD19 scFv | hN4 ΔNRR ECD | hN4 TMD | N4 JMD | G4VP 64 |
| 2A, 2C, S2C-E | CD8α SNIPR | CD8α hinge ECD with Notch1 TMD/JMD | CD19 scFv | CD8α ECD | hN1 TMD | N1 JMD | G4VP 64 |
| 2A, S2C | CD28 SNIPR | CD28 hinge ECD with Notch1 TMD/JMD | CD19 scFv | CD28 ECD | hN1 TMD | N1 JMD | G4VP 64 |

| | | | | | | | |
|---|---|---|---|---|---|---|---|
| 2A, S2C | IgG4 SNIPR | IgG4 hinge ECD with Notch1 TMD/JMD | CD19 scFv | IgG4 ECD | hN1 TMD | N1 JMD | G4VP 64 |
| S2C | OX40 SNIPR | OX40 hinge ECD with Notch1 TMD/JMD | CD19 scFv | OX40 ECD | hN1 TMD | N1 JMD | G4VP 64 |
| 2C, S2E | CD8α variant 1 | CD8α hinge ECD, variant 1, with Notch1 TMD/JMD | CD19 scFv | CD8α ECD, variant 1 | hN1 TMD | N1 JMD | G4VP 64 |
| 2C, S2E, 3B, 3D, S3A-B, S3E, 4A, 4C-F, S4A, S4E-F | CD8α variant 2 | CD8α hinge ECD, variant 2, with Notch1 TMD/JMD | CD19 scFv | CD8α ECD, variant 2 | hN1 TMD | N1 JMD | G4VP 64 |
| 2C, S2E | CD8α variant 3 | CD8α hinge ECD, variant 3, with Notch1 TMD/JMD | CD19 scFv | CD8α ECD, variant 3 | hN1 TMD | N1 JMD | G4VP 64 |
| 2C, S2E | CD8α variant 4 | CD8α hinge ECD, variant 4, with Notch1 TMD/JMD | CD19 scFv | CD8α ECD, variant 4 | hN1 TMD | N1 JMD | G4VP 64 |
| 3B | SNIPR TMD variants | hsSynNotch with variable TMD | CD19 scFv | hN1 ECD | See Table S1 | N1 JMD | G4VP 64 |
| 3B, S3A-B | Notch1 TMD alanine scan | Alanine scan of Optimized CD8α hinge Notch | CD19 scFv | CD8α ECD, variant 2 | hN1 TMD with indicated alanine mutant | N1 JMD | G4VP 64 |
| 3B, S3A-B, 4A, | CD8α variant 2, G318A | CD8α hinge ECD, variant 2, with Notch1 | CD19 scFv | CD8α ECD, variant 2 | hN1 TMD_ G318A | N1 JMD | G4VP 64 |

| | | | | | | | |
|---|---|---|---|---|---|---|---|
| 4E-F, S4F | | TMD_G318A/JMD | | | | | |
| 3B, S3A-B, 4A, 4E-F, S4F | CD8α variant 2, V319A | CD8α hinge ECD, variant 2, with Notch1 TMD_V319A/JMD | CD19 scFv | CD8α ECD, variant 2 | hN1 TMD_V319A | N1 JMD | G4VP64 |
| 3C | SNIPR JMD variants | hsSynNotch with variable JMD | CD19 scFv | hN1 ECD | hN1 TMD | See Table S1 | G4VP64 |
| 3D, S3E | CLSTN2/CLSTN2 | CD8α hinge ECD, variant 2, with CLSTN2 TMD/JMD | CD19 scFv | CD8α ECD, variant 2 | CLSTN2 TMD | CLSTN2 JMD | G4VP64 |
| 3D, S3E | CLSTN2/Notch1 | CD8α hinge ECD, variant 2, with CLSTN2 TMD/Notch1 JMD | CD19 scFv | CD8α ECD, variant 2 | CLSTN2 TMD | N1 JMD | G4VP64 |
| 3D, S3E | CLSTN2/KCNE3 | CD8α hinge ECD, variant 2, with CLSTN2 TMD/AGER JMD | CD19 scFv | CD8α ECD, variant 2 | CLSTN2 TMD | AGER JMD | G4VP64 |
| 3D, S3E | CLSTN2/PTPRF | CD8α hinge ECD, variant 2, with CLSTN2 TMD/PTPRF JMD | CD19 scFv | CD8α ECD, variant 2 | CLSTN2 TMD | PTPRF JMD | G4VP64 |
| S3C | synRobo_GV | Robo1 Fn-III domain ECD with Notch1 TMD/JMD | CD19 scFv | Fn-III ECD | Robo1 TMD_GV | N1 JMD | G4VP64 |
| S3C | HingeAGER | CD8α hinge ECD, variant 2, with AGER TMD/Notch1 JMD | CD19 scFv | CD8α ECD, variant 2 | AGER TMD | N1 JMD | G4VP64 |
| S3C | HingeAGER_LVS | CD8α hinge ECD, variant 2, with AGER TMD_LVS/Notch1 JMD | CD19 scFv | CD8α ECD, variant 2 | AGER TMD_LVS | N1 JMD | G4VP64 |

| | | | | | | | |
|---|---|---|---|---|---|---|---|
| S3D | hsNotch1 LNR, Notch2 JMD | human SynNotch1 with Notch2 JMD | CD19 scFv | hN1 ECD | hN1 TMD | N2 JMD | G4VP 64 |
| 4A, S4A | TMD Selection | CD8α hinge ECD, variant 2, with multiple TMDs/Notch1 JMD | CD19 scFv | CD8α ECD, variant 2 | See Table S1 | N1 JMD | G4VP 64 |
| 4A, S4A | JMD Selection | CD8α hinge ECD, variant 2, with multiple TMDs/Notch1 JMD | CD19 scFv | CD8α ECD, variant 2 | hN1 TMD | See Tabl e S1 | G4VP 64 |
| 4A, 4E-F, S4A-B, S4F, 5D-E, S5A, S5C-J | Hinge Notch SNIPR (Notch2 JMD) | CD8α hinge ECD, variant 2, with Notch1 TMD/Notch2 JMD | CD19 scFv | CD8α ECD, variant 2 | hN1 TMD | N2 JMD | G4VP 64 |
| 4C-D, S4D | ALPPL2 synNotch | murine SynNotch with anti-ALPPL2 scFv | ALPPL2 scFv | mN1 ECD | mN1 TMD | N1 JMD | G4VP 64 |
| 4C-D, S4D | ALPPL2 Hinge Notch SNIPR | CD8α hinge ECD, variant 2, with Notch1 TMD/JMD and anti-ALPPL2 scFv | ALPPL2 scFv | CD8α ECD, variant 2 | hN1 TMD | N1 JMD | G4VP 64 |
| 4C-D, S4D | ALPPL2 Hinge Notch SNIPR (Notch2 JMD) | CD8α hinge ECD, variant 2, with Notch1 TMD/Notch2 JMD and anti-ALPPL2 scFv | ALPPL2 scFv | CD8α ECD, variant 2 | hN1 TMD | N2 JMD | G4VP 64 |
| 4A, 4E-F, S4A, S4F | Hinge Notch SNIPR (NRG1 JMD) | CD8α hinge ECD, variant 2, with Notch1 TMD/NRG1 JMD | CD19 scFv | CD8α ECD, variant 2 | hN1 TMD | NR G1 JMD | G4VP 64 |

| | | | | | | | |
|---|---|---|---|---|---|---|---|
| S4G | Hinge Notch SNIPR (LaG17) | CD8α hinge ECD, variant 2 against eGFP | LaG17 nanobody | CD8α ECD, variant 2 | hN1 TMD | N2 JMD | G4VP64 |
| S4G | Hinge Notch SNIPR (EGFRviii scFv) | CD8α hinge ECD, variant 2 against EGFRviii | EGFRviii scFv | CD8α ECD, variant 2 | hN1 TMD | N2 JMD | G4VP64 |
| S4G | Hinge Notch SNIPR (HER2 scFv) | CD8α hinge ECD, variant 2 against HER2 | HER2 scFv | CD8α ECD, variant 2 | hN1 TMD | N2 JMD | G4VP64 |
| 5B | Humanized Hinge Notch SNIPR (Pax6) | CD8α hinge ECD, variant 2, with Notch1 TMD/Notch2 JMD | CD19 scFv | CD8α ECD, variant 2 | hN1 TMD | N2 JMD | Pax6 DBD-p65(428-551) |
| 5B, 5C-E, S5A, S5C-D | Humanized Hinge Notch SNIPR (HNF1A) | CD8α hinge ECD, variant 2, with Notch1 TMD/Notch2 JMD | CD19 scFv | CD8α ECD, variant 2 | hN1 TMD | N2 JMD | HNF1A DBD-TCR-p65(361-551) |
| 5B | Humanized Hinge Notch SNIPR (ZF6) | CD8α hinge ECD, variant 2, with Notch1 TMD/Notch2 JMD | CD19 scFv | CD8α ECD, variant 2 | hN1 TMD | N2 JMD | ZF6 - TCR - p65(361-551) |
| 5B, 5C | Humanized Hinge Notch SNIPR (ZF10) | CD8α hinge ECD, variant 2, with Notch1 TMD/Notch2 JMD | CD19 scFv | CD8α ECD, variant 2 | hN1 TMD | N2 JMD | ZF10 - TCR - p65(361-551) |

### Supplemental Figure S1

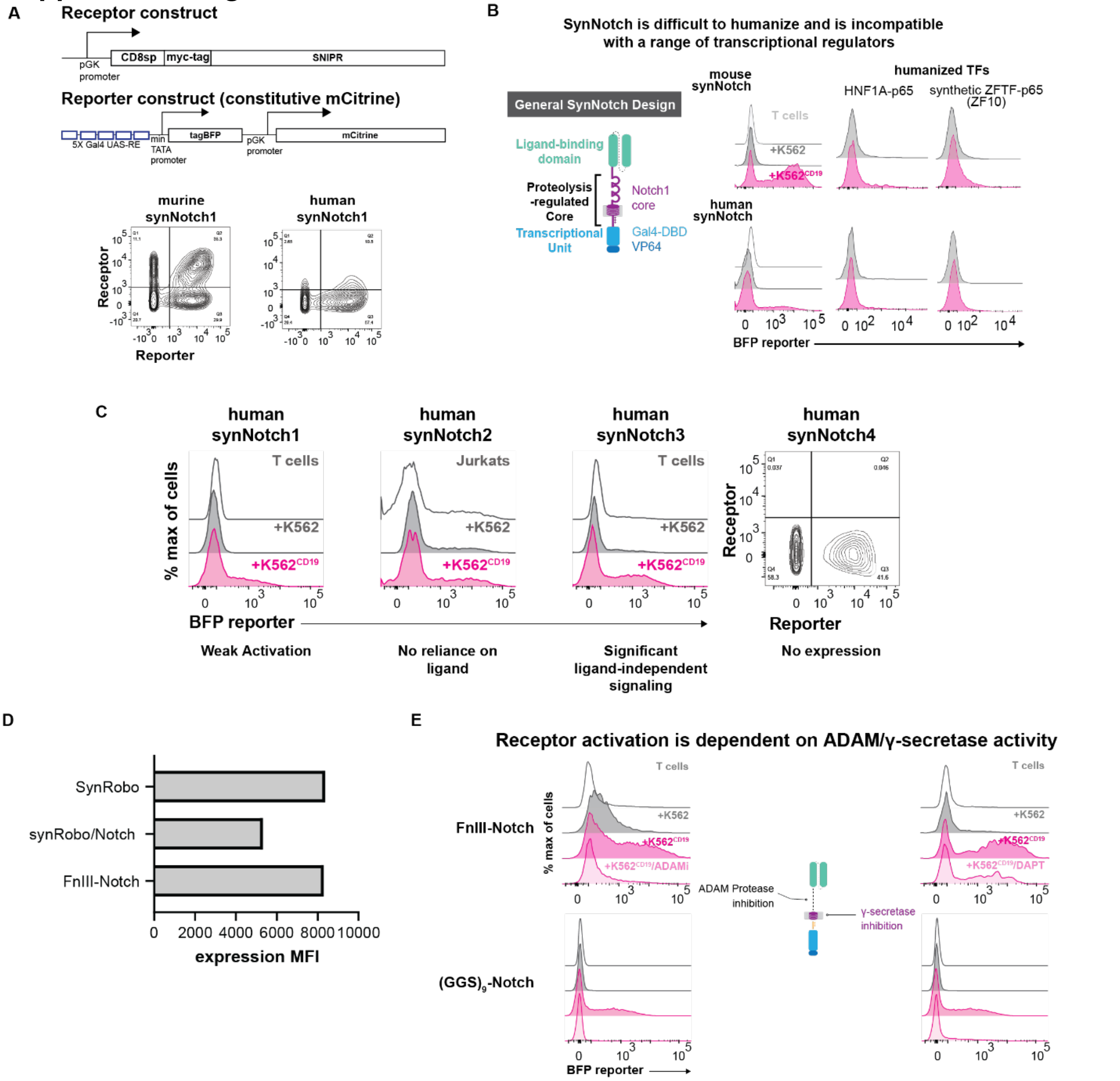


**Supplemental Figure 1. Construct designs, expression of Robo/FnIII receptors, and drug inhibition studies. (A)** Schematic of SNIPRs and reporter constructs. All SNIPRs tested are expressed under a constitutive pGK promoter and contain an N-terminal CD8α signal peptide for membrane trafficking and myc-tag for measuring expression. Expression of tagBFP is regulated by a 5x Gal4 UAS response element and a minimal pybTATA promoter. For identifying reporter+ cells, mCitrine is expressed constitutively under a pGK promoter. Right, receptor expression of murine and human synNotch receptors. **(B)** The original synNotch receptor is composed of a ligand-binding domain (LBD) fused to a proteolysis-regulated core from murine Notch1 and a Gal4-VP64 transcriptional unit. T cells expressing a CD19 synNotch receptor, constructed with either a murine or human Notch1 core, were tested with various transcriptional factors for their ability to transmit ligand-dependent signaling. SynNotch T cells were co-incubated with either K562 or K562$^{CD19}$ sender cells for 24 hours and BFP output was measured using flow cytometry proteins. Compared to the original murine synNotch design, synNotch receptors utilizing human components fail to efficiently induce BFP. (**C**) SynNotch receptors using cores from human Notch 1, Notch 2, and Notch 3 do not display strong ligand-dependent activation. Human synNotch 4 did not express. **(D)** Representative expression MFIs for synNotch/synRobo. Expression MFI does not correlate with receptor signaling ability. **(E)** Receptor activity is dependent on ADAM/γ-secretase activity. SNIPR activation is inhibited by the ADAM inhibitor GI254023X and the gamma secretase inhibitor DAPT.

### Supplemental Figure S2

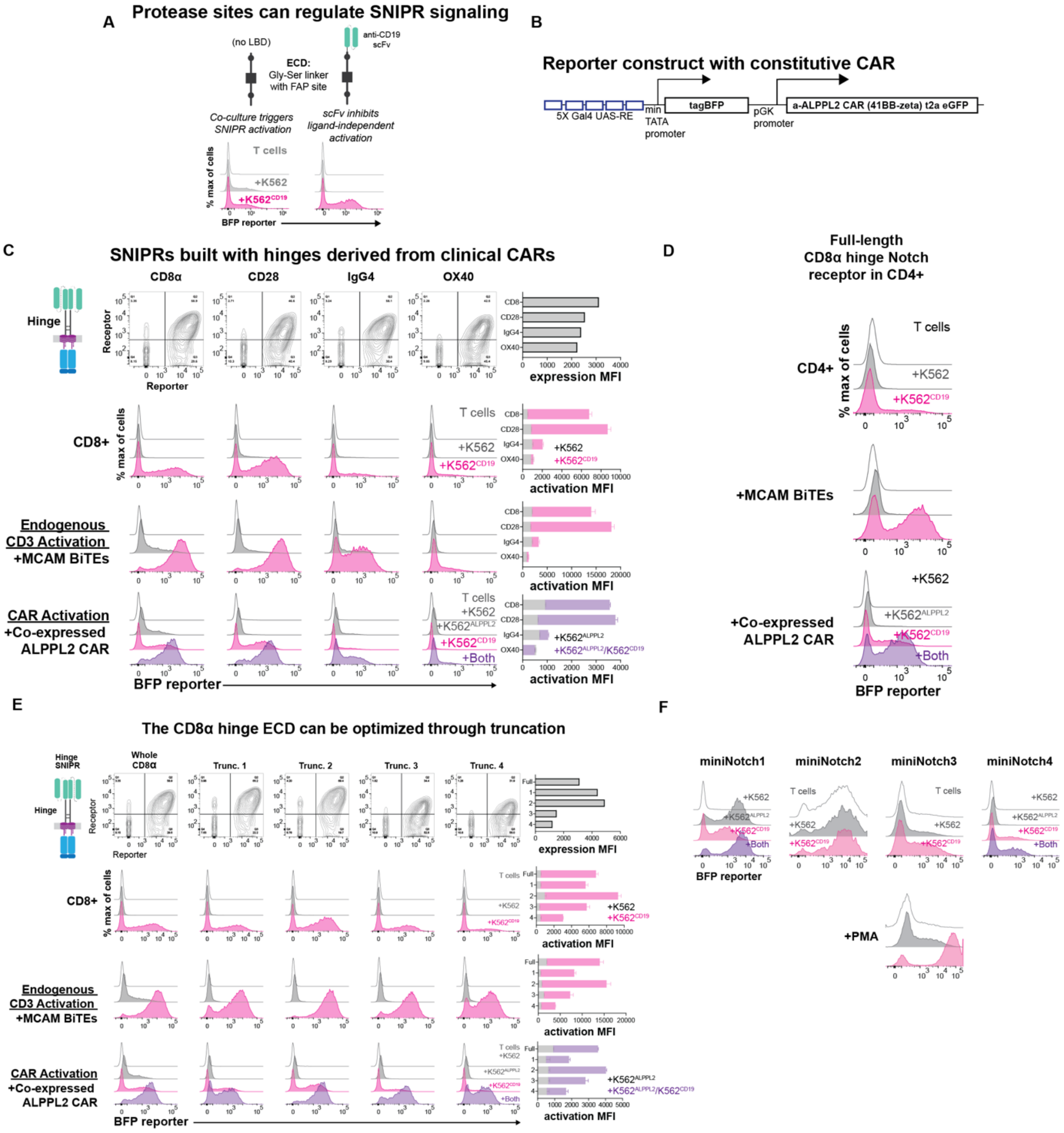


**Supplemental Figure 2. Protease site regulation, reporter constructs with CAR, hinge engineering. (A)** Protease site exposure can regulate SNIPR signaling. Addition of a fibroblast activation protein (FAP) protease site into a synthetic linker ECD confers SNIPR activity in the presence of K562 sender cells. Addition of a ligand binding domain restores ligand specificity. **(B)** Diagram of reporter construct with constitutively co-expressed 2nd generation ALPPL2 CAR (41BB-zeta). **(C)** Activation of anti-CD19 CD8α Hinge SNIPR variants. 1st row: expression of SNIPR and reporter construct in CD8+ T cells. 2nd row: Activation of the CD8α Hinge SNIPR variants with $K562^{CD19}$. 3rd row: Activation of the CD8α Hinge SNIPR variants with $K562^{CD19}$ in the presence of MCAM

BiTEs. 4th row: Activation of the CD8α Hinge SNIPR variants with $K562^{CD19}$ in the presence of a co-expressed 2nd generation ALPPL2 CAR. Superimposed bar graphs displaying ligand-independent (K562) and ligand-dependent ($K562^{CD19}$) activation are shown. **(D)** Same as C, but with the full length CD8α Hinge Notch1 expressed in CD4+ T cells. The CD8α-based SNIPR is not as sensitive to T cell activation in CD4+ T cells as in CD8+ cells. **(E)** Same as B, but with CD8α Hinge SNIPR variants.

### Supplemental Figure S3

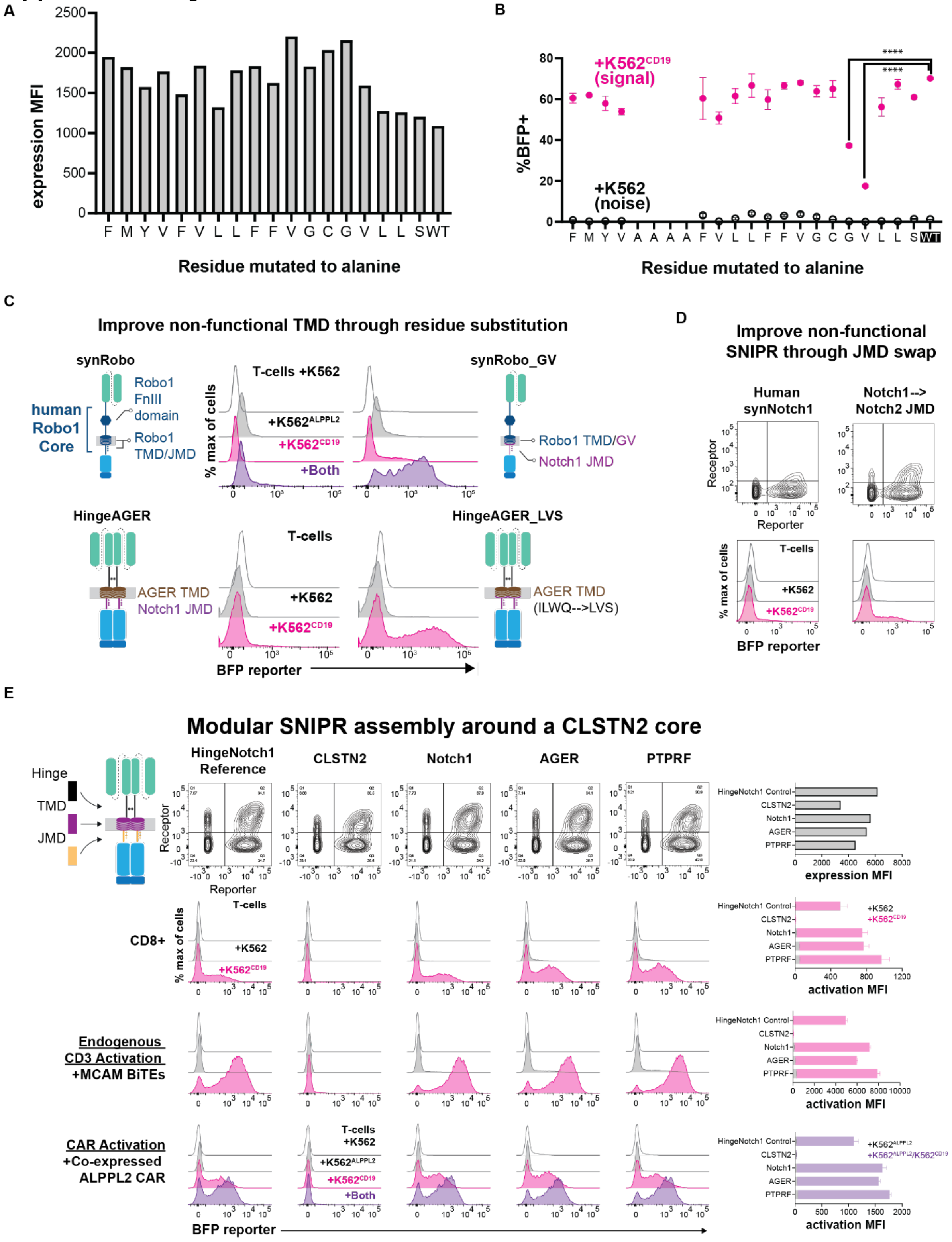

**Supplemental Figure 3. Alanine scan, fixing non-functional TMDs, non-Notch receptors. (A)** Expression of Notch1 TMD Alanine scan mutants. **(B)** Results from Alanine scan in terms of %BFP+. **(C)** Fixing a non-functional TMD through residue substitution. The performance of the Robo1 TMD in a SNIPR setting can be improved by inserting a Gly-Val motif at an equivalent position to the Notch1 TMD and replacing the Robo1 JMD with that of Notch1. The performance of the AGER TMD in a SNIPR setting can be improved by replacing the residues C-terminal to its Gly-Val motif with those from the highest performing TMD, Notch1 from *G. gallus*. T cells expressing a SNIPR-BFP circuit were co-incubated with the indicated K562 cells for 48 hours and the percentage of BFP+ cells was measured using flow cytometry. (**D**) Human synNotch expression and activation improvement through JMD substitution identified by a screen. Replacing the human Notch 1 JMD with the Notch 2 JMD increases receptor expression and activation in primary T cells. **(E)** Activation of anti-CD19 SNIPRs with non-Notch components. $1^{st}$ row: expression of SNIPR and reporter construct in CD8+ T cells. $2^{nd}$ row: Activation of the SNIPR variants with $K562^{CD19}$. $3^{rd}$ row: Activation of the CD8α Hinge SNIPR variants in the presence of MCAM BiTEs. $4^{th}$ row: Activation of the CD8α Hinge SNIPR variants in the presence of a co-expressed $2^{nd}$ generation anti-ALPPL2 CAR. Superimposed bars displaying ligand-independent (K562) and ligand-dependent ($K562^{CD19}$) activation are shown.

### Supplemental Figure S4

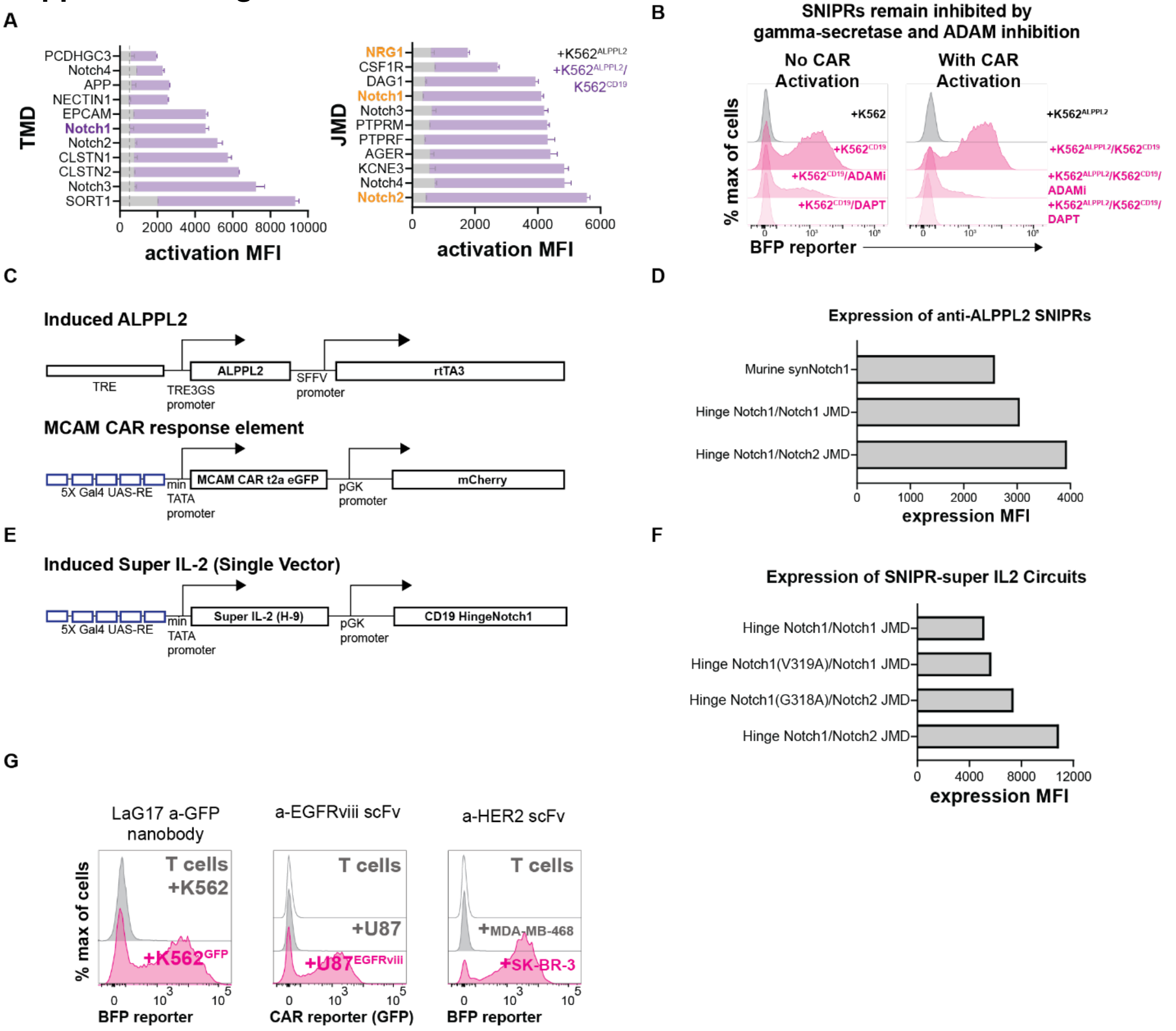


**Supplemental Figure 4. Hinge SNIPR TMD and JMD, drug inhibition studies, construct designs, expression data, testing with additional LBDs. (A)** Superimposed bar graphs displaying activation of Hinge SNIPRs with variable TMDs and JMDs with an activated co-expressed ALPPL2 CAR at 48 hours. **(B)** SNIPR activation is dependent on ADAM and gamma-secretase activity. T cells expressing a CD19 Hinge Notch SNIPR-BFP circuit and co-expressed ALPPL2 CAR were co-incubated with the indicated conditions for 48 hours. BFP output was measured using flow cytometry. **(C)** Design of inducible ALPPL2 cassette and MCAM CAR response element. rtTA3 is expressed under an SFFV promoter and induces ALPPL2 expression in the presence of doxycycline. MCAM CAR is expressed under an inducible minimal TATA promoter and 5XGal4 UAS enhancer with a constitutively expressed mCherry. **(D)** Expression of ALPPL2 SNIPR was measured using myc-staining. **(E)** Design of an induced Super IL-2 single vector. Super IL-2 is expressed under an inducible minimal TATA promoter and 5XGal4 UAS enhancer with a constitutively expressed SNIPR. **(F)** Expression of SNIPR-super IL-2 circuits were measured using myc-staining. **(G)** Activation of Hinge SNIPR with additional LBDs. Hinge SNIPR with Notch2 JMD is effective against membrane bound GFP, EGFRviii, and HER2 when expressing an anti-GFP nanobody, anti-EGFRviii scFv, or anti-HER2 scFv, respectively.

### Supplemental Figure S5

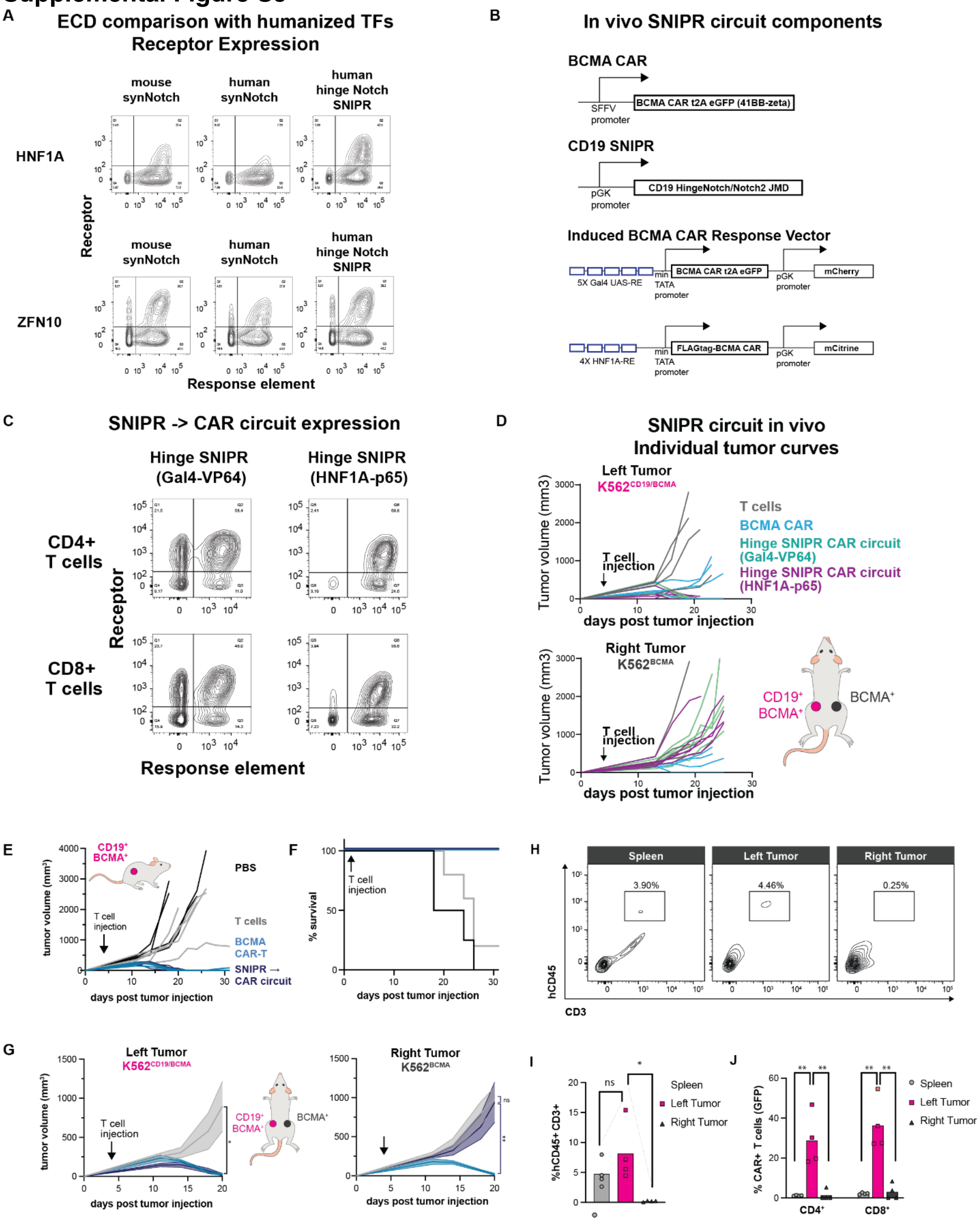


**Supplemental Figure 5. Constructs, expression of humanized and non-humanized receptors, and individual tumor growth curves. (A)** Comparison of humanized TFs with SNIPR ECDs. Expression profile of optimized CD8α Hinge SNIPRs and synNotch receptors using the human transcription factor HNF1A and synTF ZFN10 was determined by surface staining of the myc-tagged receptors. **(B)** Design of BCMA CAR, CD19 SNIPR, and induced BCMA CAR response vector. BCMA CAR is expressed under a constitutive SFFV promoter. CD19 SNIPR is expressed under a constitutive pGK promoter. BCMA CAR is expressed under an inducible minimal TATA promoter and 5XGal4 UAS enhancer with a constitutively expressed mCherry. **(C)** Receptor expression of optimized CD8α Hinge SNIPRs in CD4+ and CD8+ T cells used for *in vivo* testing of circuit function. **(D)** Individual tumor curves for the averaged data in Fig 5E. (**E**) Gal4 SNIPR-CAR circuits selectively eliminate tumors with dual antigen signatures in vivo. NOD mice (5 per experimental group) were injected with $1x10^6$ $K562^{CD19+/BCMA+}$ target cells into the left flank. 4 days post tumor injection, $5x10^6$ BCMA CAR or CD19 SNIPR-BCMA CAR circuit T cells ($2.5x10^6$ each CD4+ and CD8+) or PBS control were injected, and tumor size was measured over time. **(F)** Survival curve for mice in E. **(G)** Same as A, but with NOD mice injected with $1x10^6$ $K562^{CD19+/BCMA+}$ target cells into the left flank and $1x10^6$ $K562^{BCMA}$ target cells into the right flank. **(H)** Human T cell presence in the spleen, left, and right tumors was measured using flow cytometry. **(I, J)** CAR induction of human T cells in the spleen, left, and right tumors was measured using a t2a system. Statistics were calculated using one-way ANOVA with Dunnett's test post hoc. $^*P \leq 0.05$; $^{**}P \leq 0.01$; ns, not significant.
